# A comprehensive physiological dataset of pain and aversive modalities for pain biomarkers research

**DOI:** 10.64898/2026.09.29.755497

**Authors:** Alyson Champagne, Lorie-Ève Barrette, Antoine Cyr-Bouchard, Mathieu Roy, Michel-Pierre Coll

**Affiliations:** School of Psychology, Université Laval, Québec, QC, Canada; Centre interdisciplinaire de recherche en réadaptation et intégration sociale, Centre intégré universitaire de santé et de services sociaux (CIUSSS) de la Capitale-Nationale, Québec, QC, Canada; Department of Psychology, McGill University, Montréal, QC, Canada

**Author notes:** Correspondence Assistant Professor, École de psychologie, Pavillon Félix-Antoine-Savard, 2325, rue des Bibliothèques Université Laval, Québec (Québec) G1V 0A6.

## Abstract

While progress has been made in identifying pain biomarkers using neuroimaging, the development of reliable EEG-based biomarkers remains challenging, limited by small sample sizes and the lack of adequate control conditions. We present a comprehensive dataset from 125 pain-free adults, comprising electroencephalogram (EEG) recordings, and behavioural, sociodemographic, and psychological data. Galvanic skin response and electrocardiogram data were also recorded in a subset of participants (*n* = 76). Participants received two sustained painful stimulations (thermal, chemical) and two sustained aversive non-painful stimulations (auditory, taste), each delivered twice (with and without continuous intensity ratings). Data are provided in Brain Imaging Data Structure format, where all physiological, stimulus-related, and behavioural data are synchronized within the EEG data files for easy access and analysis. Technical validation confirmed data integrity, with spectral and physiological analyses showing responses associated with stimulus intensity and subjective pain/unpleasantness levels. By providing multimodal physiological data across painful and non-painful aversive conditions, this initiative represents a valuable resource for developing or validating biomarkers and exploring the multidimensional nature of pain perception.

## Background & Summary

Pain is a complex and subjective experience influenced by multiple aspects, such as contextual, affective and cognitive factors^1,2^. Considerable efforts have been devoted to identifying reliable brain-based biomarkers, ranging from diagnostic and prognostic tools for chronic pain to biomarkers intended to predict treatment response or quantify pain intensity^3,4^. Considering its ease of use, low cost, and good reliability^5^, electroencephalography (EEG) has been widely used in the search for pain biomarkers^6^. EEG studies have shown that pain perception is associated with changes in neural oscillations, such as suppression of alpha-band activity^7–11^, both increases and decreases in beta-band power^12–14^, and enhancement of gamma-band activity^7,10,11,14,15^. Beyond cerebral activity, pain triggers responses from the autonomic nervous system, such as galvanic skin response (GSR) and changes in heart rate (HR), which both generally show a peak following a noxious stimulus, before gradually returning to baseline^16–20^. Moreover, sociodemographic (e.g., age, sex) and psychological characteristics (e.g., anxiety, depression, pain catastrophization) can further influence how pain is perceived^21,22^.

Recently, there has been a growing interest in applying machine learning algorithms to decode or predict perceived pain or pain intensity from neurophysiological data. These methods can leverage the multivariate nature of such signals to identify complex pain-related patterns^3,6,23,24^. Research on autonomic signals has also demonstrated that physiological responses provide valuable information for decoding pain^25^. However, several methodological limitations have hindered the development of robust EEG-based pain biomarkers, such as small sample sizes (median of 24 participants in ^6^) and insufficient external validation across independent datasets resulting in limited generalizability^3,4,6^. Furthermore, assessing the specificity of developed biomarkers remains challenging since many studies rely on a single modality to induce pain (e.g., thermal stimulation) and do not include appropriate control conditions to distinguish pain-specific responses from those reflecting general stimulus salience or negative valence^26,27^.

The scientific community is addressing these limitations by releasing more datasets publicly. For instance, Zhao *et al.* obtained EEG recordings and pain ratings from 678 healthy participants across eight studies involving phasic laser stimulations^28^. Similarly, Jones *et al.* recorded EEG, HR, oxygenation, and behavioural data from 112 newborns during a painful heel lance, a sham, or a non-noxious auditory condition^29^. The *PainMonit* Dataset provides nine physiological signals from 55 healthy adults receiving painful heat stimulations and from 49 participants assessing pain during a physiotherapy session^30^. Chowdhury *et al.* also made available EEG and behavioural data from 150 healthy adults experiencing prolonged experimental temporomandibular pain^31^. Data from many other EEG studies delivering pain are openly available^32–35^, as well as standardized analysis pipelines to facilitate preprocessing, analysis and visualization of EEG data, and reproducibility^36^. Despite these advancements in data sharing, most existing open-access datasets focus on a single modality to induce pain and do not offer comprehensive non-painful aversive controls. To address these gaps, we therefore provide a dataset that combines multimodal physiological signals with diverse experimental conditions, offering a comprehensive framework for developing and validating robust and pain-specific biomarkers.

The initial release of our dataset includes a comprehensive set of variables from 125 pain-free adults. EEG recordings, physiological data (*n* = 76; GSR and electrocardiogram (ECG)), and continuous ratings are provided during two tonic pain stimulations (heat of varying intensities and orofacial pain from capsaicin-rich hot sauce) and two aversive stimulations (pure tones of varying intensities and bitter melon paste). EEG and physiological recordings are also available during two resting-state conditions (eyes open and closed), along with self-reported sociodemographic and psychological characteristics (anxiety, depression, and pain catastrophization) for each participant. To ensure data integrity, we performed a thorough technical validation. We first preprocessed the EEG data using an adaptation of the DISCOVER-EEG pipeline^36^ in Python. We then analyzed EEG spectral power across sensors and frequency bins based on subjective ratings (painful vs non-painful, unpleasant vs not unpleasant) and on objective stimulus intensity for thermal pain and aversive auditory stimulations. For the physiological data, we extracted standardized metrics of skin conductance and HR to verify their quality and characterize autonomic reactivity across stimulus intensities, as well as based on subjective ratings. These analyses confirmed that both neurophysiological and autonomic responses capture biologically plausible modulations across experimental conditions. Additionally, preliminary analyses using part of our data showed that thermal pain and matched auditory stimulations could be discriminated using the EEG signals within and across individuals^37^. Ultimately, by providing diverse high-quality physiological data obtained from both painful and non-painful aversive conditions, this initiative aims to facilitate collaborative efforts toward more reproducible, generalizable, and nuanced models of pain perception.

## Methods

### Participants

Data from 125 pain-free adult participants are available. Sociodemographic and psychological characteristics are presented in Table 1. Participants were recruited through email advertisements sent to students and staff at a Université Laval in Québec, Canada, as well as through posters displayed at *Université Laval* and at the Centre for Interdisciplinary Research in Rehabilitation and Social Integration (Cirris, Québec, Canada). All participants reported not meeting any of the following exclusion criteria: (a) experiencing chronic or persistent pain, (b) being pregnant or breastfeeding, (c) having any neurologic or psychiatric diagnosis, (d) taking medication that alters attention or vigilance, and (e) not having normal or corrected-to-normal vision, audition, and olfaction. The experimental session lasted three hours and was conducted at the Cirris. All participants gave their written informed consent for their participation and anonymized data sharing, and received 45 CAD for their participation.

**Table 1.**
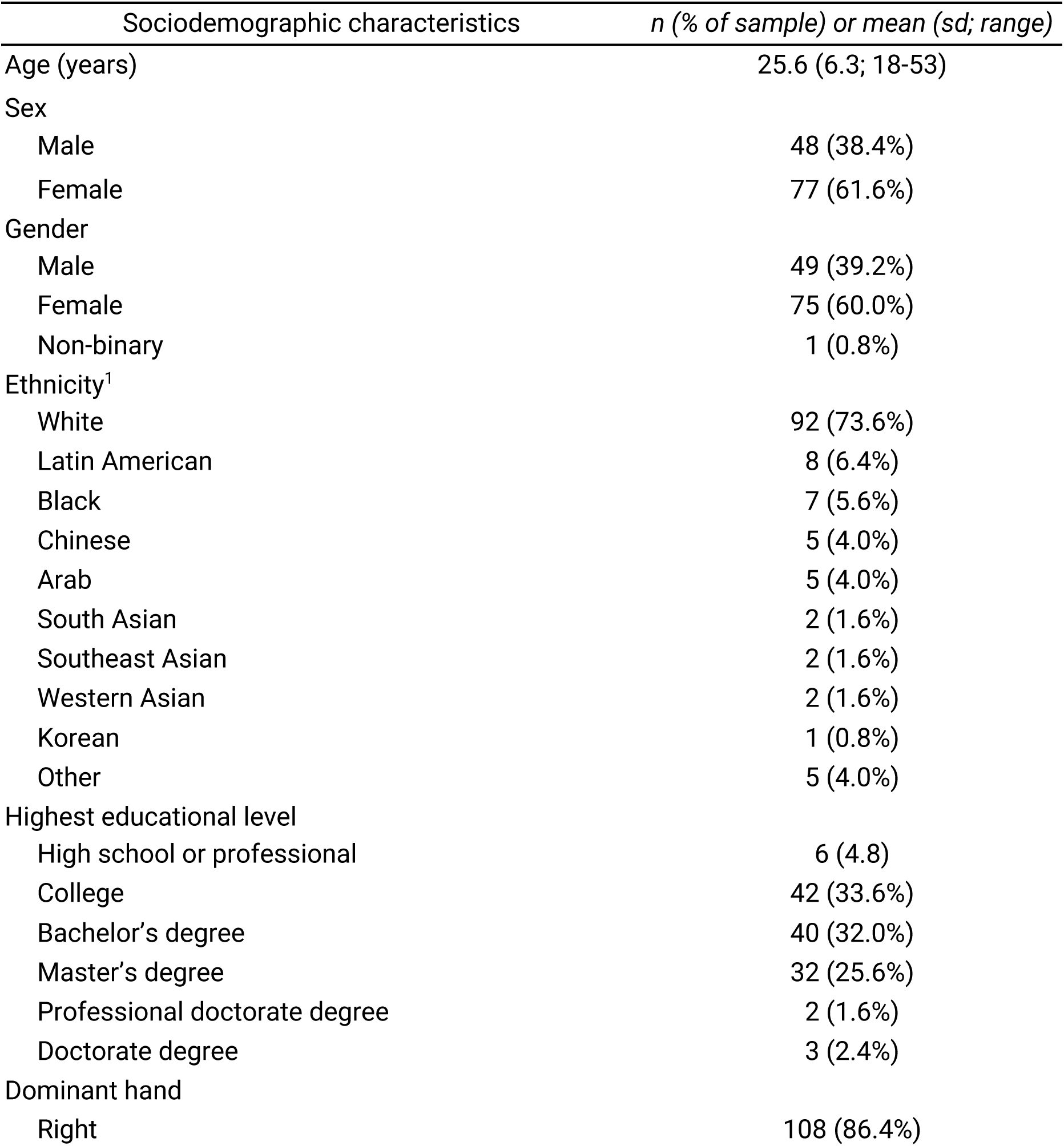
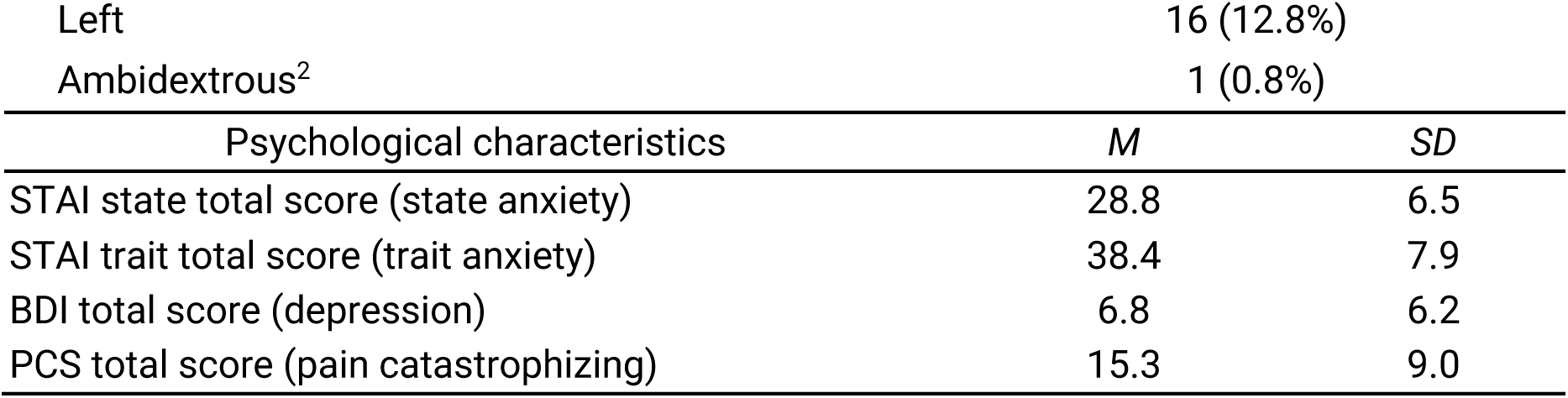
Sociodemographic and psychological characteristics. *n* = 125. *STAI* = State-Trait Anxiety Inventory, *BDI* = Beck Depression Inventory, *PCS* = Pain Catastrophizing Scale. ^1^Since participants could select more than one ethnicity, the total value might differ from 100. ^2^For participants who reported being ambidextrous, the hand preferred for writing was designated as the dominant hand for the purpose of this study.

| Sociodemographic characteristics | *n* (% of sample) or mean (sd; range) |
| --- | --- |
| Age (years) | 25.6 (6.3; 18-53) |
| Sex |  |
| Male | 48 (38.4%) |
| Female | 77 (61.6%) |
| Gender |  |
| Male | 49 (39.2%) |
| Female | 75 (60.0%) |
| Non-binary | 1 (0.8%) |
| Ethnicity<sup>1</sup> |  |
| White | 92 (73.6%) |
| Latin American | 8 (6.4%) |
| Black | 7 (5.6%) |
| Chinese | 5 (4.0%) |
| Arab | 5 (4.0%) |
| South Asian | 2 (1.6%) |
| Southeast Asian | 2 (1.6%) |
| Western Asian | 2 (1.6%) |
| Korean | 1 (0.8%) |
| Other | 5 (4.0%) |
| Highest educational level |  |
| High school or professional | 6 (4.8) |
| College | 42 (33.6%) |
| Bachelor's degree | 40 (32.0%) |
| Master's degree | 32 (25.6%) |
| Professional doctorate degree | 2 (1.6%) |
| Doctorate degree | 3 (2.4%) |
| Dominant hand |  |
| Right | 108 (86.4%) |

**Table 1. Sociodemographic and psychological characteristics.**
| Left | 16 (12.8%) |  |
| --- | --- | --- |
| Ambidextrous<sup>2</sup> | 1 (0.8%) |  |
| Psychological characteristics | *M* | *SD* |
| STAI state total score (state anxiety) | 28.8 | 6.5 |
| STAI trait total score (trait anxiety) | 38.4 | 7.9 |
| BDI total score (depression) | 6.8 | 6.2 |
| PCS total score (pain catastrophizing) | 15.3 | 9.0 |

This study was approved by the Sectorial Ethics Committee for Rehabilitation and Social Integration Research of the *Centre intégré universitaire de santé et de services sociaux de la Capitale-Nationale* (#2023-2487).

### Experimental design

The design followed a three-step sequence: (1) individualized calibration for the aversive auditory and the thermal pain modalities, (2) resting-state EEG recordings, and (3) delivery of four types of stimulation, either painful (thermal, chemical) or aversive (auditory, taste) with and without continuous ratings of pain or unpleasantness (see Figure 1).

**Figure 1.**
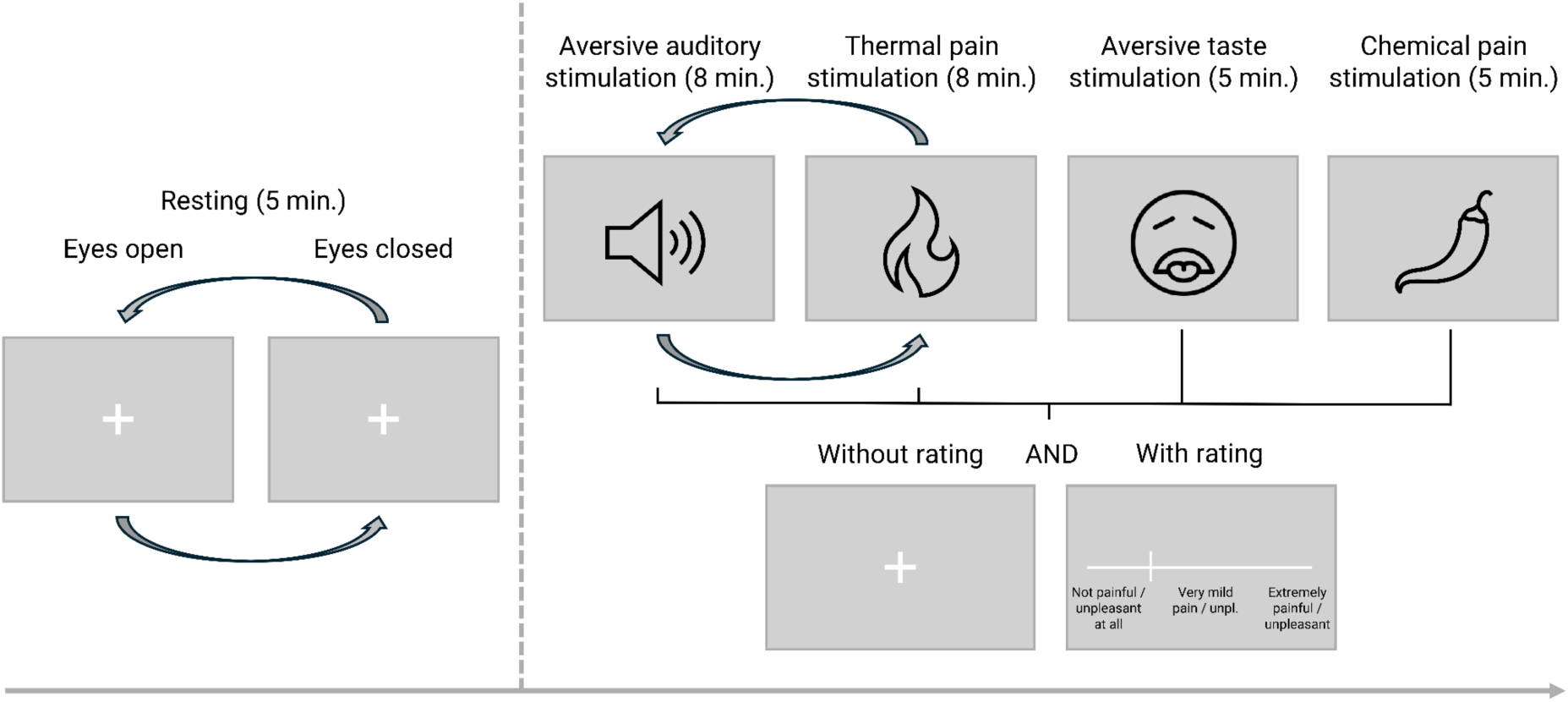
Design of the EEG experiment after thermal and auditory calibration tasks. Participants first completed two resting-state recordings (eyes-open and eyes-closed). They then completed four pain or aversive stimulation tasks: aversive auditory (8 min.), thermal pain (8 min.), aversive taste (5 min.), and chemical pain (5 min.). Arrows represent counterbalanced tasks. Aversive taste and chemical pain tasks were always administered last to prevent carryover effects. Each task was administered twice: once without and once with continuous ratings using response buttons to move a cursor on a visual analogue scale. EEG was recorded continuously throughout the session. Unpl. : unpleasantness.

*Tonic thermal pain stimulation.* Thermal pain stimuli were delivered using two different research thermal stimulators due to an equipment update during the data collection period: a TSA-II (Medoc Ltd, Israel; *n* = 18) equipped with a 3.00 x 3.00 cm ATS probe, and a TCS-II (QST Lab, France; *n* = 107) equipped with a T09 probe featuring five 0.74 x 1.22 cm thermodes.

Stimulation temperatures were individually calibrated based on each participant’s perception using a task implemented in *PsychoPy* (v2024.2.4), an open-source software package for stimulus presentation and experimental control^38^. Five target temperatures (40 °C, 42 °C, 44 °C, 46 °C, and 48 °C) were each administered four times on the volar surface of the non-dominant forearm, with a duration of 20 s per trial. Stimuli were presented in a fixed pseudorandomized order, ensuring that no temperature was repeated consecutively. The stimulation site on the forearm was systematically shifted between four different zones across trials to minimize sensitization effects and to maintain perceptual sensitivity throughout the calibration task. To avoid skin burns and exceeding the devices built-in safety parameters during the eight-minute thermal stimulations in the EEG experiment (see *EEG experiment*), 48 °C was chosen as the maximal temperature applied.

During thermal calibration, participants continuously rated their perceived pain on a visual analogue scale (VAS) ranging from “No pain” (*Aucune douleur*) to “Extremely painful” (*Extrêmement douloureux*) by moving a cursor represented by a small white rectangle using two response buttons. A label indicating “Very mild pain” (*Très faible douleur*) was also placed at the midpoint of the scale. If a temperature became intolerable at any point, the thermode was removed from their arm and participants were instructed to place the cursor at the “Extremely painful” mark on the VAS.

The cursor position was subsequently converted into a numerical score ranging from 0 to 200 (0 = “No pain”, 100 = “Very mild pain”, 200 = “Extremely painful”). This rating procedure was identical for all calibration and experimental tasks. Ratings and timestamps were recorded at a frequency of 50 Hz, allowing a detailed analysis of participants’ perception. Individual calibration data were then modeled to estimate the temperatures corresponding to three target pain levels: hot, but non-painful (VAS-50), mildly painful (VAS-100; pain detection threshold), and moderately painful yet still tolerable (VAS-150; see dashed line in Figure 2). Specifically, for each participant, maximum ratings across all administered temperatures were extracted and fitted with linear, hyperbolic, quadratic, and exponential functions. The resulting best-fitting model was determined with a least-squares optimization approach and used to interpolate the temperatures corresponding to the target ratings. These temperatures were then imputed in a custom Python script to create a 490 s file used for the EEG experiment with the thermode. In a subset of participants (*n* = 48), target temperatures were manually adjusted to comply with the devices’ safety limits. Each intensity was presented three times in a predetermined order, with 4 s ramps ensuring gradual transitions between conditions (see dashed line in Figure 2). Between each temperature, the thermode was pseudo-randomly moved to one of four possible positions on the arm to minimize sensitization. A baseline temperature of 38 °C was applied before and after each stimulation, during which the temperatures corresponding to VAS-50, VAS-100, and VAS-150 intensities were delivered for 64 s, 54 s, and 44 s, respectively. These durations were chosen to evoke a tonic stimulation eliciting different levels of pain and non-painful thermal stimuli, while minimizing habituation. Across participants, temperatures corresponding to VAS-50, VAS-100, and VAS-150 intensities were on average 43.0 °C (± 1.8 °C), 45.2 °C (± 1.6 °C), and 47.0 °C (± 1.2 °C), respectively.

**Figure 2.**
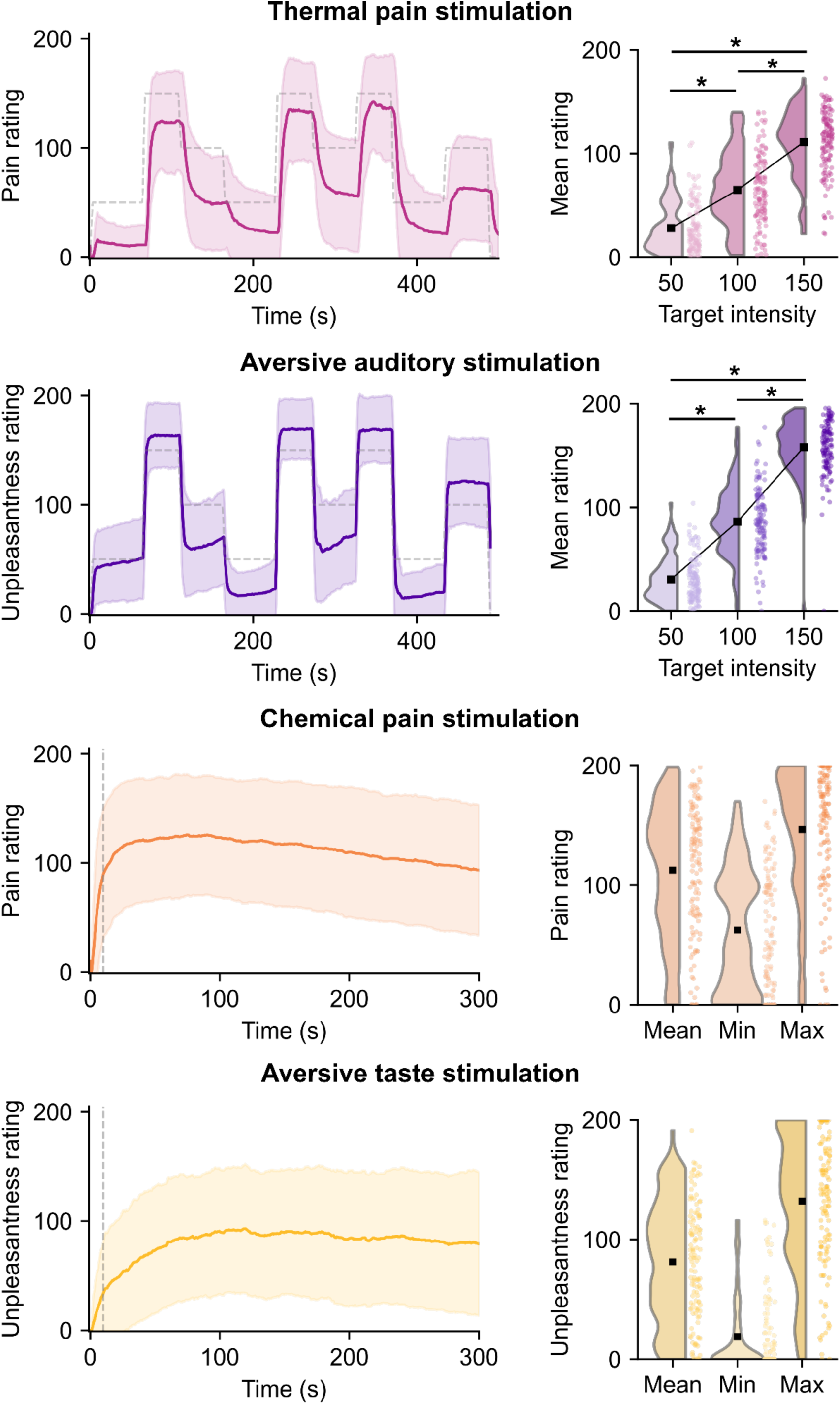
Average subjective pain and unpleasantness ratings across the four stimulations. Left-side panels show the average subjective ratings over time. The solid colored line represents the mean rating across all participants, while the shaded area represents the standard deviation (SD), calculated within each 0.2 s time point, indicating inter-participant variability. For the thermal pain and aversive auditory tasks, dashed grey lines indicate the objective intensity of the stimulus where applicable. For the chemical pain and aversive taste stimulations, the first 10 s were excluded from subsequent EEG and physiological analyses due to increased movement at stimulus onset (represented by the vertical dashed line here). Right-side panels show the mean subjective ratings grouped by target intensity (thermal and auditory) or the mean, minimum, and maximum subjective ratings by condition (chemical and taste). Data are illustrated as raincloud plots, where the shaded violin plots represent the distribution of ratings and individual dots represent mean ratings for each participant. The black squares represent the group means. Horizontal black bars and asterisks (*) indicate statistically significant differences between intensities (*p* < .001). Unpl.: unpleasantness; 50: VAS-50 target intensity; 100: VAS-100 target intensity; 150: VAS-150 target intensity.

*Aversive auditory stimulation.* This modality was selected as a non-painful aversive control for the tonic thermal pain stimulation, as it allowed for a similar temporal manipulation of stimulation intensities, facilitating the matching of salience across sensory modalities. A 1,000 Hz pure-tone sound delivered through a standard pair of headphones (*OneOdio Pro 10*) connected to a computer was used to induce varying levels of aversive auditory stimulation. This frequency was chosen as it offers a wider range to modulate intensity and discomfort according to individual thresholds. While frequencies between 2,000 and 4,000 Hz are generally considered the most aversive^39^, they more easily reach intolerable or unsafe levels, limiting the flexibility needed for the calibration of multiple stimulation intensities^40^.

Auditory stimulation was calibrated using a custom *PsychoPy* script. The 1,000 Hz pure-tone sound was presented at five different decibel (dB) levels: 50 dB, 60 dB, 70 dB, 80 dB and 90 dB, with a sampling frequency of 44,100 Hz. *PsychoPy* generates sounds using linear amplitude scaling rather than absolute dB sound pressure level (SPL) values. To ensure that each stimulus corresponded to the intended SPL, a single, independent acoustic verification was performed prior to the start of data collection using a calibrated sound level meter and a reference tone. This procedure established the mapping between linear amplitude values in *PsychoPy* and actual dB SPL values. All auditory stimuli lasted 20 s and were presented binaurally via headphones. Each intensity level was presented four times across 20 trials to ensure robust sampling of subjective unpleasantness elicited by each sound. The sequence of stimulus presentation was identical across all participants, following a predetermined order of varying intensities where both lower- and higher-intensity tones were interspersed throughout the calibration. No intensity level was presented twice in consecutive trials. This order was designed to prevent rapid adaptation to a single intensity and to maintain participants’ sensitivity throughout the calibration task.

During the auditory calibration, participants were asked to continuously rate their level of discomfort on a VAS ranging from “Not unpleasant at all” (*Aucunement désagréable*) to “Extremely unpleasant” (*Extrêmement désagréable*) by moving a cursor represented by a small white rectangle using two response buttons. A label indicating “Very mild unpleasantness” (*Très faible désagrément*) was also placed at the midpoint of the scale. If a sound was intolerable, participants were instructed to remove the headphones and to place the cursor at the “Extremely unpleasant” mark on the VAS. Such intensities were excluded from the subsequent EEG experiment.

The procedure for determining the three target auditory intensities (VAS-50; loud, but not unpleasant, VAS-100; mildly unpleasant, and VAS-150; moderately unpleasant) for the EEG experiment was similar to the one used for thermal pain stimulation. The cursor position was converted into a numerical score, and ratings and their corresponding timestamps were recorded at a frequency of 50 Hz. A custom Python script generated an individualized audio stimulus file (.wav) based on the calibration procedure. Four mathematical models (linear, hyperbolic, parabolic, and exponential) were fitted to each participant’s calibration data using a least-squares optimization approach. The resulting best-fitting model was then used to derive the specific sound intensities (dB) associated with each target rating. These values were subsequently converted into linear amplitude units to ensure compatibility with *PsychoPy*’s sound generation framework. Finally, the script synthesized a 490 s audio file in which each intensity was presented three times in a fixed order (see dashed line in Figure 2). During each presentation, the VAS-50, VAS-100, and VAS-150 tones lasted 64, 54, and 44 s, respectively, including smooth 4 s amplitude ramps implemented between conditions to ensure gradual transitions. Across participants, the sound intensities corresponding to VAS-50, VAS-100, and VAS-150 stimulation levels were on average 61.5 ± 8.9 dB, 71.4 ± 8.9 dB, and 80.3 ± 7.5 dB, respectively.

*Tonic chemical pain stimulation.* Orofacial chemical pain was induced using Tabasco Habanero sauce (7,000 Scoville units; www.tabasco.com/hot-sauces/habanero-sauce; as in ^41^). A 0.1 mL dose of capsaicin-rich hot sauce was placed on an edible filter paper and spread over a 1-cm circle. Participants then placed the side of the filter paper with the sauce on their tongue and kept their mouth closed for 30 s. Afterward, the paper was removed, and participants breathed through their mouth for one minute to allow the sauce to dry at a specific point on the tongue. They then closed their mouth for the duration of each EEG experiment (five minutes). This procedure was repeated twice, once before each task (see *EEG experiment*).

*Aversive taste stimulation.* This stimulation served as an aversive control for the orofacial chemical pain task, as both stimuli shared the same spatial location and a similar temporal profile, with both sensations peaking early and slowly subsiding over the five-minute recording period. A small amount (0.5 g; 1/8 teaspoon) of bitter melon powder (*Lily of the Valley Organic Bitter Melon (Karela) powder*) was mixed with an equal amount of water to create a paste, which was then placed on an edible filter paper. This substance was selected as it is known for its intense bitter flavor, which is generally perceived as unpleasant while being safe for consumption. Participants then placed the paper in their mouth for the entire duration of the EEG experiment, which was five minutes. This procedure was repeated twice, once before each task (see *EEG experiment*).

*Resting-state tasks.* The brain’s electrical signals were first recorded during two five-minute resting-state tasks: one with eyes open and one with eyes closed. The eyes-open task was initially prioritized to ensure methodological consistency with the stimulation tasks, which required visual fixation or evaluation. However, since many studies use a closed eye resting-state task^42–44^ and that brain oscillatory activity differs significantly between these states^45^, an eyes-closed task was later introduced in the study (*n* = 78 completed only the eyes–open condition, and *n* = 47 completed both the eyes–open and eyes–closed conditions). During both tasks, participants were instructed to relax and limit their movements. For eyes-open condition, they were instructed to fixate on a white cross displayed at the center of the screen with a gray background to reduce ocular artifacts.

*EEG experiment.* A schematic representation of the experiment is shown in Figure 1. All experimental tasks were programmed and administered using *PsychoPy*. The presentation order of the resting-state tasks was randomized, as was the order of the aversive auditory and thermal pain stimulations, across participants. Specifically, two possible sequences were used for the experimental tasks: (1) aversive auditory stimulations (eight minutes), painful thermal stimulations (eight minutes), aversive taste stimulations (five minutes), and painful chemical stimulations (five minutes), or (2) painful thermal stimulations, aversive auditory stimulations, aversive taste stimulations, and painful chemical stimulations. The aversive taste and chemical pain tasks were systematically administered last to prevent residual oral sensations from contaminating other modalities. Furthermore, the chemical pain task always concluded the session to ensure that participants never received two different painful stimulations (thermal and chemical) consecutively.

EEG recordings was collected during the application of the four stimulations, which were each administered twice: first in a passive task (during which the participant received the stimulation while looking at a white cross in the middle of the screen) and then in an active task (the participant continuously rated the painfulness or unpleasantness of the stimuli using response buttons to move the cursor on the VAS). This procedure allowed for the measurement of stimulation intensity both with and without the associated motor response, as well as the cognitive processes involved in the explicit evaluation of pain and unpleasantness. In the active task, participants responded using the index and middle fingers of their dominant hand.

### Data acquisition

*EEG recordings and physiological metrics.* Brain electrical activity was recorded in a sound-attenuated booth using Acticap Slim electrodes (Brain Products, Germany) and amplified with a NA300 amplifier (EGI, USA, *n* = 49, 63 electrodes) connected through an actiCAP ControlBox and an adapter interface, or an actiCHamp Plus amplifier (Brain Products, Germany, *n* = 76, 63 electrodes). These changes resulted from an equipment upgrade that occurred during the course of the study. The signals remained comparable as the same electrodes were used. Data were acquired at a sampling rate of 1000 Hz with an online filter between DC and 500 or 280 Hz. The electrodes were placed following the international 10-10 system^46^, and their impedance was kept below 10 kΩ throughout recordings. The ground electrode was placed at position FPz, and the reference electrode at position FCz. Table 2 shows the number of participants with available EEG and behavioural data for each task. Data were missing either because participants could not tolerate the stimulation and did not complete the task, or because poor signal quality led to their exclusion. Additionally, ratings were missing for one participant during the thermal pain stimulation task due to a technical error.

**Table 2.** EEG, physiological (GSR and ECG), and behavioural data available and usable for each task. Missing EEG data occurred for some participants due to intolerable stimulations. For the physiological data, eight participants had noisy or flat GSR and ECG recordings. The resting-state task with eyes closed, as well as the physiological recordings, were introduced later in the study, resulting in a smaller number of available data.

| Task | *n* (EEG) | *n* (physio) |
| --- | --- | --- |
| Aversive auditory stimulation without rating | 124 | 69 |
| Aversive auditory stimulation with rating | 124 | 69 |
| Thermal pain stimulation without rating | 124 | 68 |
| Thermal pain stimulation with rating | 123 | 68 |
| Aversive taste stimulation without rating | 125 | 69 |
| Aversive taste stimulation with rating | 124 | 69 |
| Chemical pain stimulation without rating | 125 | 69 |
| Chemical pain stimulation with rating | 123 | 67 |
| Resting-state task with eyes opened | 125 | 69 |
| Resting-state task with eyes closed | 47 | 45 |

For participants with the actiCHamp Plus amplifier, horizontal electrooculography (HEOG) was recorded using two electrodes placed approximately 1 cm lateral to the outer canthus of each eye. Vertical electrooculography (VEOG) was recorded from the left eye only, with two electrodes positioned above the eyebrow and under the eye. GSR was collected using the Brain Products GSR module (Brain Products, Germany). Two electrodes were placed on the middle phalanx of the index and middle fingers of the non-dominant hand (15 mm x 36 mm x 36 mm), and an isotonic electrolyte gel (*EDALYT*) mimicking the concentration of salt in sweat was applied. GSR was recorded using the exosomatic approach with direct current, where a constant voltage of 0.5 V was applied as recommended^47^. ECG recordings were acquired using a BIP2AUX adapter (Brain Products, Germany) in a Lead 1 configuration, with electrodes positioned on the right collarbone (ground), the left collarbone (anode), and the left floating ribs (cathode).

*Statistical analysis of behavioural data for technical validation*. To validate our experimental manipulations, subjective ratings were analyzed for each of the four stimulation modalities. We first made sure that all rating values remained within the valid 0 to 200 numerical range on the VAS. For the thermal pain and aversive auditory stimulations, we then computed mean ratings for each participant across the three target intensity levels (VAS-50, VAS-100, and VAS-150) by averaging the rating values within the duration of each trial (each target intensity was presented three times, see *Experiment design*). Since the chemical pain and aversive taste tasks involved tonic stimulations with an initial peak in intensity, participants tended to exhibit increased movement at the onset of the stimulation. To minimize movement-related artifacts in subsequent behavioral, EEG and physiological analyses, the first 10 s of the task were excluded. Overall mean, minimum and maximum ratings were then calculated. Time spent above a rating of 100 on the VAS, associated with the presence of pain or unpleasantness, was also computed.

Statistical analyses were performed using the Pingouin package (v0.5.5)^48^ in Python. To assess the effect of thermal and auditory stimulus intensity on subjective perception, a one-way repeated-measures ANOVA was conducted with the target intensity as the within-subject factor. If significant main effects were detected, post-hoc pairwise *t*-tests were performed using Bonferroni correction for multiple comparisons.

*EEG data preprocessing.* To ensure EEG data standardization, facilitate sharing and enhance reproducibility, all raw data was first converted into the Brain Imaging Data Structure (BIDS) format^49,50^. Information regarding acquisition parameters, channel definitions, event markers, and other details were written into the BIDS structure.

EEG data preprocessing was performed using a custom pipeline built around the MNE-BIDS (v0.16.0)^50,51^ and MNE-Python (v1.9.0)^52^ packages, and adapted from the DISCOVER-EEG pipeline^36^. First, a 60 Hz line noise was removed from the continuous data using a notch filter to eliminate power-line artifacts, followed by a 0.5 Hz high-pass filter, a 100 Hz low-pass filter, and resampling to 500 Hz. Faulty EEG channels were detected and marked for each participant using the pyprep package (v0.4.3)^53,54^ if they showed consistently low correlation with neighboring channels (*r* < 0.4) or if their z-scored amplitude deviated substantially from the median channel amplitude (*z* > 5). Channels remaining after this process were considered “good” channels. The data was then re-referenced to the common average, and segmented into 5 s epochs with a 50% overlap. Subsequent artifact rejection was performed at the epoch level. To correct for physiological artifacts, Independent Component Analysis (ICA) was conducted using the extended Infomax algorithm^55^. For a robust decomposition, the ICA was fitted on data that was high-pass filtered at 1 Hz and excluding epochs with peak-to-peak amplitudes exceeding 500 µV. The resulting components were automatically classified using the ICLabel algorithm^56^ with the MNE-ICALabel package^57^, and components labeled as “muscle artifact”, “eye blink”, “heart beat”, “line noise” or “channel noise” with a probability greater than 70% were considered artifacts and removed. The percentage of components labeled as “brain” activity out of the total ICA-derived components was computed. Transient epoch-level artifacts remaining after ICA correction were corrected using the AutoReject algorithm^58^ by cross-validating the optimal number of interpolated channels among candidate values (4, 8, or 16), allowing for a maximum interpolation of ∼25% of the montage. Epochs remaining after this step were considered “good” epochs. Finally, channels identified as faulty were interpolated using spherical splines before subsequent analyses, such as feature extraction and time-frequency decomposition to ensure the same data structure across participants.

*Statistical analysis of EEG data for technical validation and quality control.* We characterized the spectral profiles across the subjective experiences of pain or unpleasantness and the different stimulus intensities. Power spectral density (PSD) was first computed for each 5 s epoch using the multitaper method^59,60^ in MNE-Python and covering a frequency range of 1 to 100 Hz. Epochs were zero-padded to 10 s prior to PSD estimation to achieve a frequency resolution of 0.1 Hz. Moreover, to account for inter-individual variability and to provide a standardized measure of spectral changes, we normalized PSD data for each task relative to the participant’s resting-state baseline (eyes open) using a dB transform: (*10* ∗ *log10*(*Task*/*Rest*)), which addresses the 1/*f* distribution of EEG power and ensures that increases and decreases relative to baseline are represented symmetrically^61^. To further characterize spectral changes independently of the aperiodic component, power spectra were parametrized using the specparam algorithm, formerly known as *Fitting Oscillations & One-Over-F* (FOOOF)^62^, applied to each participant’s mean power spectrum per condition for each EEG channel over 1 to 100 Hz, with the following settings, based on ^62^: peak width limits = [1, 6] Hz, maximum number of peaks = 6, minimum peak height = 0.05, peak threshold = 1.5, fixed aperiodic mode.

To characterize the electrophysiological modulations associated with subjective experience and stimulus intensity, statistical analyses were performed on the periodic component of the power spectrum derived from the specparam decomposition described above. Epochs rated as painful or unpleasant (mean epoch VAS rating ≥ 100) were compared to those rated as not painful or not unpleasant (mean epoch VAS rating < 100) using *t*-tests. We also explored the effect of stimulus intensity (VAS-50, VAS-100, VAS-150) for the thermal pain and aversive auditory tasks using repeated-measures ANOVAs. Statistical significance was determined using non-parametric cluster-based permutation tests to handle the multiple comparisons across EEG sensors and frequency bands^63^. Clusters were defined as groups of adjacent sensors and frequency bands, and frequency bins exceeding a cluster-entering threshold of *p* < .01, two-tailed. The significance of these clusters was then determined using 5,000 permutations with a cluster-level significance threshold of *p* < .05, incorporating a “hat” correction to the *t*-test statistic^64^. These tests were applied to: (1) grand-average topographical maps for standard frequency bands (Theta: 4-8 Hz; Alpha: 8-13 Hz; Beta: 13-30 Hz; Gamma: 30-80 Hz), using a spatial adjacency matrix to identify significant electrode clusters in each frequency band, and (2) global periodic PSD averaged across channels to identify significant frequency clusters.

*Physiological data preprocessing.* Physiological signals (GSR and ECG) were processed using the NeuroKit2 toolbox (v0.2.12)^65^ in Python. Data quality was first assessed through visual inspection of the raw GSR and ECG signals. Participants with very noisy or flat recordings in both GSR and ECG channels were excluded from physiological analyses (*n* = 8).

GSR data were first converted to microsiemens. The continuous recording for each task was then cleaned using the BioSPPy algorithm^66^ and downsampled to 10 Hz. The signal was decomposed into tonic (skin conductance level; SCL) and phasic (SCR) components using the cvxEDA convex optimization approach^67^, implemented in Neurokit2. Only the phasic component was retained for subsequent analyses. For subjective experience analyses across all tasks, GSR signals were segmented into 20 s epochs categorized based on the mean VAS rating during the epoch. Since participants were exposed to ongoing stimulations, non-specific SCR frequency (SCRs per minute) was computed, as these responses indicate overall electrodermal activity over time rather than reflecting a single evoked response^47,68^. For intensity analyses of the aversive auditory and thermal pain stimulations, GSR signals were segmented into nine 40 s epochs time-locked to the onset of changes in stimulus intensity. This length was chosen to capture the full electrodermal response and recovery period, as SCR half-recovery generally ranges from 2 to 10 s while the overall response cycle can extend beyond 20 s^47^. The magnitude of the first SCR exceeding 0.01 μS^47,69^ whose onset occurred within a 5-8 s time window following stimulus onset was extracted since these analyses involve varying, identifiable intensity changes. This time window skips the stimulus ramp (4 s after stimulation onset), takes into account the minimum SCR latency of 1 s, and lasts the typical SCR onset latency of 1-4 s after stimulus change^47,69^. Epochs with no detectable SCR were scored as 0 μS, while those excluded due to signal-quality failure were treated as missing data.

Raw ECG signals were first filtered with a band-pass Finite Impulse Response (FIR) filter (0.5 to 40 Hz)^70^. QRS complexes were localized based on the steepness of the absolute gradient of the ECG signal using the Neurokit2 algorithm. R-peaks were then detected as local maxima within the QRS complexes and subsequently corrected for artifacts (e.g., missed, extra, or misaligned beats) using the automated beat-correction algorithm implemented in Neurokit2^71^. For each epoch, we extracted the weighted mean HR relative to the eyes-open resting-state baseline (ΔHR, in beat per minute (bpm)). Both the epoch-level HR and the resting-state baseline were computed using the weighted mean heart period method, which corrects for the sampling bias introduced by averaging HR over a variable number of beats^70,72^. To ensure the plausibility of our ECG data, we only retained epochs with a mean HR between 40 and 120 bpm. Epochs for the subjective experience analyses lasted 20 s, with categorization based on the mean VAS rating across this time window. For the thermal pain and aversive auditory tasks, the ECG-derived HR metric was computed over a 4-24 s window post-onset to exclude the transient cardiac orienting response associated with the onset of the intensity ramp. For the chemical and aversive taste tasks, which lack an intensity ramp, the first 10 s of the recording were excluded (as noted above) before segmenting the remainder into consecutive 20 s epochs. For intensity analyses, epochs were time-locked to the onset of each intensity plateau (VAS-50, VAS-100, and VAS-150), with HR computed over the 4-24 s window.

*Statistical analysis of physiological metrics for technical validation and quality control.* Statistical analyses of physiological metrics were performed using linear mixed-effects models (LMM) with the statsmodels package (v0.14.4)^73^ in Python. To assess the effects of subjective experience, epochs were categorized as painful/unpleasant (mean epoch VAS rating ≥ 100) or non-painful/non-unpleasant (mean epoch VAS rating < 100) and entered as a categorical fixed factor. For stimulus intensity analyses for both thermal pain and aversive auditory tasks, the target stimulus level (VAS-50, VAS-100, VAS-150) was mean-centered on VAS-100, rescaled to unit steps of 50, and treated as a continuous fixed effect. The resulting coefficient thus reflects the change associated with one step in target level intensity. In both analyses, individual epochs were entered as separate observations rather than averaged within condition or intensity level, allowing the model to capture trial-to-trial variability in addition to inter-individual variability. Prior to analysis, SCR metrics were log(1 + x)-transformed to address the positive skew and the presence of zero-amplitude epochs in electrodermal data^69^. All physiological metrics were then Z-score standardized across the whole sample. For each analysis, we specified a maximal model with a random intercept and a random slope per subject. Models that failed to converge were simplified to a structure including only a random intercept. All models were estimated using maximum likelihood and statistical significance was set at *p* < .05. To control for multiple comparisons, *p*-values were corrected using the Benjamin-Hochberg false discovery rate procedure^74^.

*Self-reported questionnaires.* Data from five questionnaires are available for all participants. First, a sociodemographic questionnaire was completed to document several demographic characteristics based on the 2021 Canadian census form (e.g., age, gender, education level, ethnicity). Three psychological questionnaires were also administered. The Pain Catastrophizing Scale (PCS)^75^ is a 13-item self-reported scale that measures the tendency to view pain as a significant event in daily life. The French-Canadian version has excellent internal consistency (α = 0.91) and good test-retest reliability (*r* = 0.85)^76^. The Beck Depression Inventory (BDI) assessed depressive symptoms^77^. Its French-Canadian version demonstrates good psychometric properties (internal consistency, α = 0.92, and test-retest reliability, *r* = 0.62)^78^. To measure general anxiety levels as well as present-state anxiety, the State-Trait Anxiety Inventory (STAI) was administered^79^. The French-Canadian version of the questionnaire has good construct validity, high internal consistency (α = 0.90 for state anxiety, and α = 0.91 for trait anxiety), and adequate test-retest reliability over a 14-day period (*r* = 0.64 for state anxiety, and *r* = 0.94 for trait anxiety)^80^. All questionnaires were completed at the end of the experimental session, except for the STAI state anxiety questionnaire (Y1), which was administered at the beginning of the session to assess anxiety at the time of testing, given its potential impact on pain perception^81^.

### Data Record

All raw data is accessible on OpenNeuro^82^ (https://openneuro.org/datasets/ds008349). It is presented in accordance with the BIDS standards as shown in Figure 3, and all modalities are synchronized within the EEG data files for easy access and analysis. For all tasks involving participants’ ratings (i.e., “active” tasks), the following files are provided : *sub-XXX_task-TaskName_stim.json* and *sub-XXX_task-TaskName_stim.tsv.gz*. The former contains the metadata detailing task description and technical recording specifications, while the latter includes continuous ratings recorded throughout the task in their original sampling rate. Additionally, behavioural ratings, stimulation intensity and thermode temperature were upsampled to match the EEG sampling rate, synchronized with the EEG signal, and added as miscellaneous channels within the EEG data files. These channels are also present but filled with −1 when not relevant (e.g., temperature channel for the auditory task) to ensure the same data structure across all files. Consequently, the EEG files contain all relevant experimental data, including EEG recordings, physiological signals (GSR, ECG, EOG), and stimulation/behavioural measurements.

**Figure 3.**
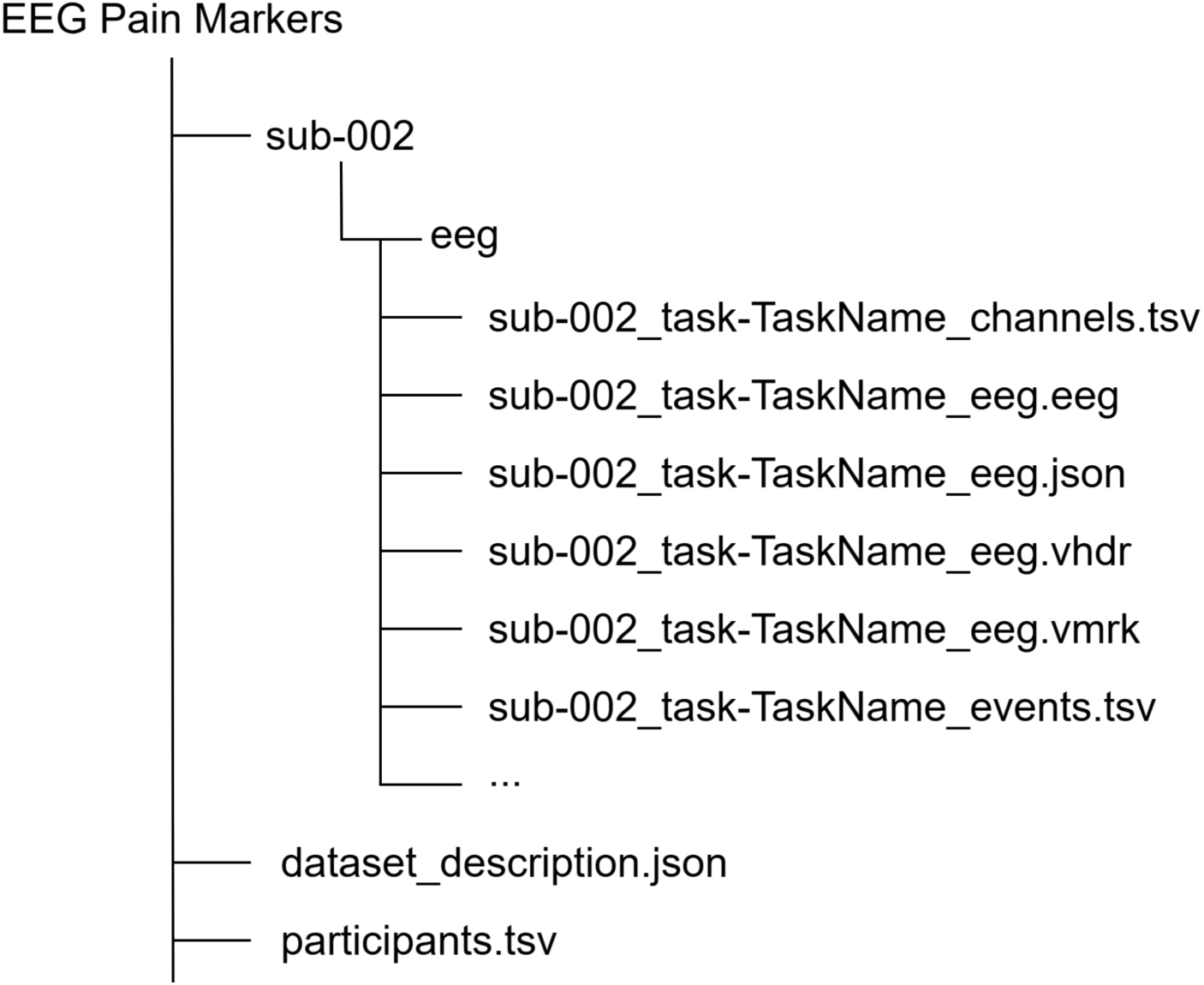
BIDS data structure. “Taskname” is a generic label representing the specific experimental conditions included in the dataset: *audioactive, audiopassive, thermalactive, thermalpassive, tasteactive, tastepassive, chemicalactive, chemicalpassive, restingopen,* and *restingclose*.

All relevant experimental information is included in the dataset. The *participants.tsv* contains all sociodemographic and psychological data acquired from the questionnaires, selected intensities following the audio and thermal calibration, as well as information about each task (e.g., duration, number and types of channels, sampling frequency). It also includes comments recorded by the experimenter for each session, originally written in French and translated into English.

### Technical validation

#### Task characteristics and evaluation

To validate the efficacy of the experimental protocol in eliciting pain or discomfort, we assessed the intensity of participant’s subjective ratings in all four tasks. The average pain or unpleasantness ratings, as well as their time course, during each active task are shown in Figure 2.

Across all active tasks, participants’ ratings on the VAS ranged from 0 to 200. For the thermal pain stimulation, mean ratings for the VAS-50, VAS-100, and VAS-150 target intensities were 28.11 ± 23.10, 64.76 ± 33.02, and 111.00 ± 30.47, respectively. The one-way repeated-measures ANOVA revealed a significant main effect of stimulus intensity on subjective pain ratings (*F*(2,244) = 531.84, *p* < .001, *η*^2^_g_ = 0.58). Post-hoc pairwise comparisons using Bonferroni correction confirmed significant differences between all intensity levels. Specifically, ratings for the VAS-100 intensity were significantly higher than those for the VAS-50 intensity (*t*(122) = 15.65, *p* < .001, *d* = 1.29). Ratings for the VAS-150 intensity were significantly higher than both the VAS-100 intensity (*t*(122) = 17.50, *p* < .001, *d* = 1.46) and the VAS-50 intensity (*t*(122) = 31.33, *p* < .001, *d* = 3.07).

For the aversive auditory task, mean ratings for the VAS-50, VAS-100, and VAS-150 target intensities were 30.62 ± 20.90, 86.08 ± 31.52, and 157.99 ± 26.07, respectively. A significant main effect of stimulus intensity on subjective unpleasantness ratings was found (*F*(2,246) = 1559.62, *p* < .001, *η*^2^_g_ = 0.80), and post-hoc pairwise comparisons also confirmed significant differences between all intensity levels. Ratings for the VAS-100 intensity were significantly higher than those for the VAS-50 intensity (*t*(123) = 25.52, *p* < .001, *d* = 2.07). Ratings for the VAS-150 intensity were significantly higher than both the VAS-100 intensity (*t*(123) = 29.75, *p* < .001, *d* = 2.49) and the VAS-50 intensity (*t*(123) = 56.27, *p* < .001, *d* = 5.39).

For the chemical pain task, the mean rating was 112.50 ± 52.75, with a mean minimum of 62.47 ± 49.76 and a mean maximum of 146.54 ± 52.83. Finally, for the aversive taste task, the mean rating was 81.43 ± 47.56, with a mean minimum of 18.76 ± 31.49 and a mean maximum of 132.23 ± 55.27.

The duration during which ratings exceeded 100 (pain/unpleasantness threshold) varied by task: participants spent on average 189 s (± 58 s; range: 0–350 s; 38% of task duration) above this threshold during aversive auditory stimulation, 137 s (± 73 s; range: 0– 460 s; 28%) during painful thermal stimulation, 110 s (± 96 s; range: 0–302 s; 37%) during aversive taste stimulation, and 171 s (± 109 s; range: 0–302 s; 57%) during painful chemical stimulation. In the aversive taste task, a majority of participants felt unpleasantness, but a higher variability was observed. In each task, some participants reported no unpleasantness or pain throughout the stimulation (auditory: *n* = 1; thermal: *n* = 4; taste: *n* = 35; chemical: *n* = 20), accounting for the lower bound of 0 s.

### EEG data preprocessing and quality control characteristics

To assess the quality of EEG data, we adapted the quality control pipeline proposed in ^83^, where (1) the percentage of “good” channels, (2) the percentage of independent components (ICs) related to brain source activity, and (3) the percentage of “good” epochs were assessed^83^. Across all tasks, participants had an average of 1.1 ± 1.7 channels marked as faulty, which represented 98.2% ± 2.8% of “good” channels. On average, 12.9 ± 4.2 “bad” ICA components were removed per participant, while 37.1% ± 10.3% of all components were identified as brain sources with a confidence score above 0.7. After epoch rejection, an average of 97.1% ± 3.0% of trials were retained for final analysis. Quality metrics per task are shown in Figure 4. Overall, these results indicate high quality EEG data. Our low channel rejection rate (∼ 2%) and epoch retention rate (∼ 97%) compare favorably to benchmarks established during the validation of the DISCOVER-EEG pipeline on other datasets^36^. For instance, validation of this pipeline on the LEMON^84^ and TDBRAIN^85^ open-access datasets reported average channel rejection rates ranging from 2% to 7% and trial retention rates between 93% and 96%. Furthermore, our average proportion of brain sources is similar to other studies^86^.

**Figure 4.**
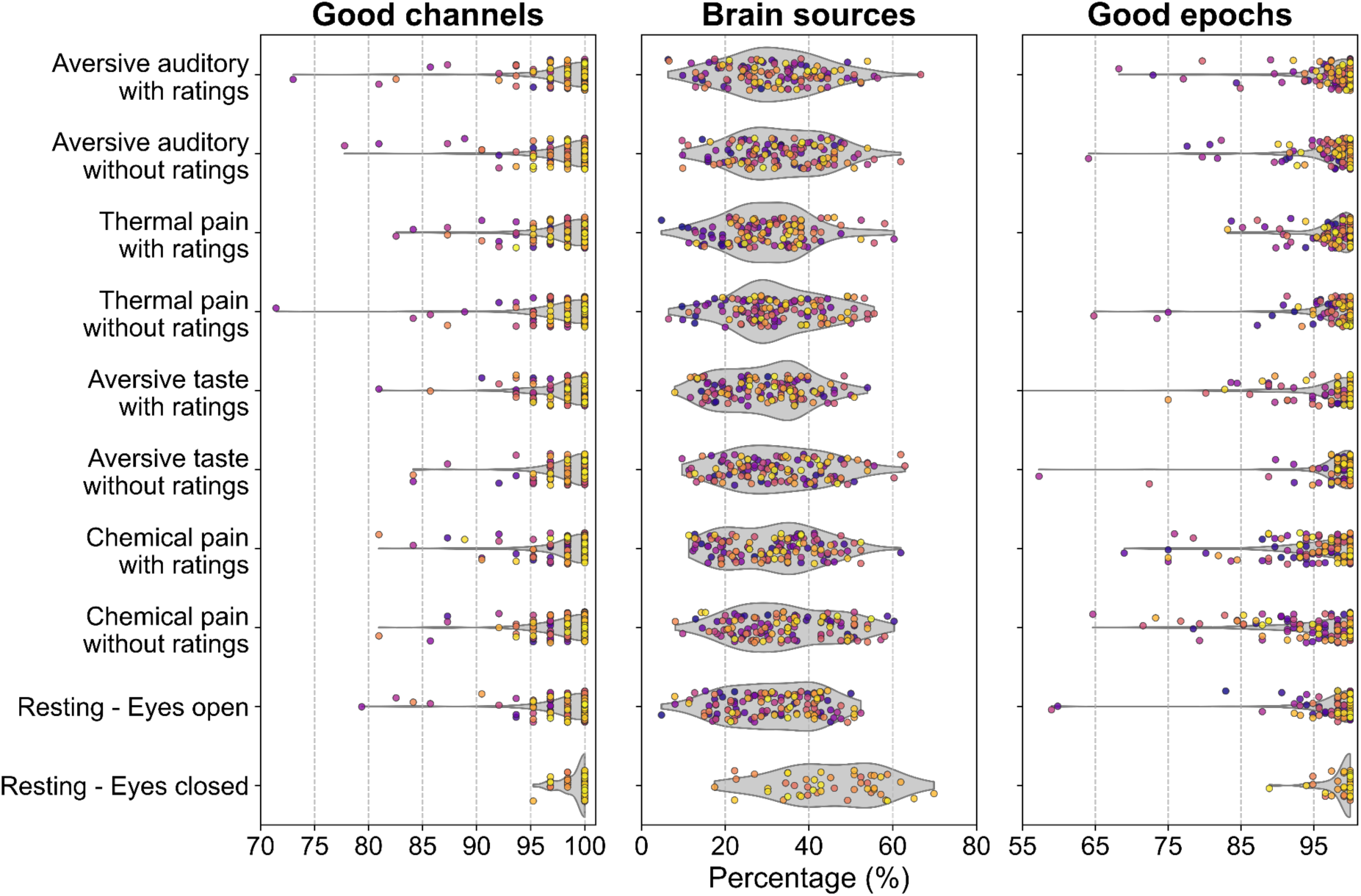
EEG quality control characteristics. Three quality metrics (i.e., the percentage of “good” channels, the percentage of ICs related to brain source activity, and the percentage of “good” epochs) were assessed across all tasks. Each participant is represented by a dot.

### Power spectral density and topographical maps

To validate the sensitivity of the EEG data to both subjective perception (Figure 5 to 8) and stimulus intensity (Figure 9 and 10), we examined the spectral and topographical distributions of neural activity across all tasks. Participants were excluded if they reported only pain or unpleasantness (taste: *n* = 6, chemical: *n* = 35) or never reported pain or unpleasantness across the whole task (auditory: *n* = 1, thermal: *n* = 4, taste: *n* = 35, chemical: *n* = 20), resulting in final samples of *n* = 123 (auditory), *n* = 119 (thermal), *n* = 81 (taste), and *n* = 58 (chemical) for these analyses.

**Figure 5.**
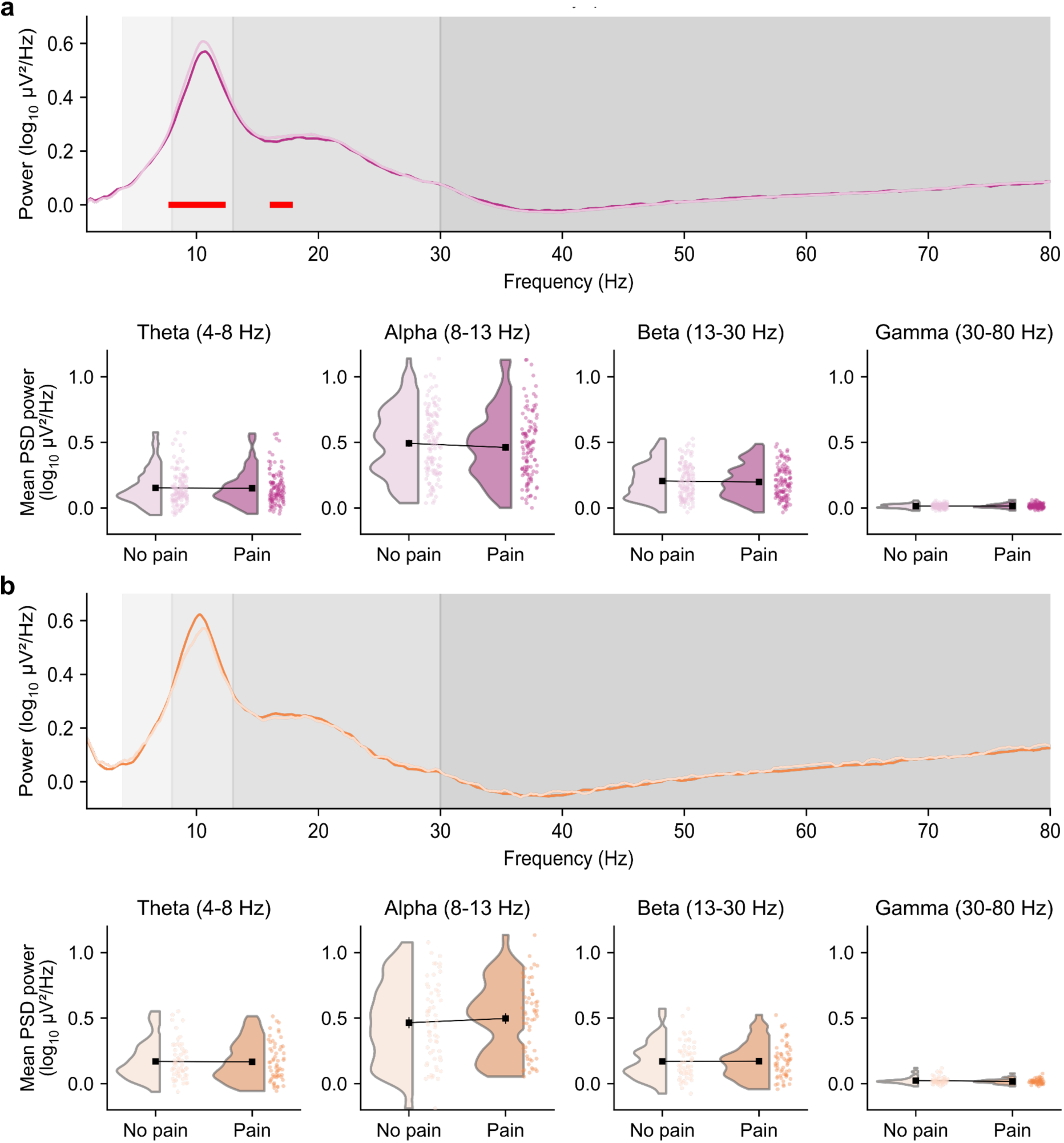
Global periodic PSD based on subjective ratings for (a) thermal and (b) chemical pain tasks. Lighter colors represent non-painful epochs (VAS ≤ 99), while darker colors represent painful epochs (VAS ≥ 100). Top panels display grand-average periodic power spectra across a frequency range of 4 to 80 Hz. Grey-shaded background regions indicate standard frequency bands: Theta (4–8 Hz), Alpha (8–13 Hz), Beta (13–30 Hz), and Gamma (30–80 Hz), respectively. Red horizontal bars show significant frequency clusters identified by the non-parametric cluster-based permutation tests (*p* < .05). The PSD between 58 to 62 Hz was interpolated to correct for the notch filter artifact. Bottom panels show the mean periodic PSD power by frequency band grouped by condition (no pain vs pain). Data are illustrated as raincloud plots, where the shaded violin plots represent the distribution of PSD power and individual dots represent mean PSD power for each participant. The black squares represent the group means.

**Figure 6.**
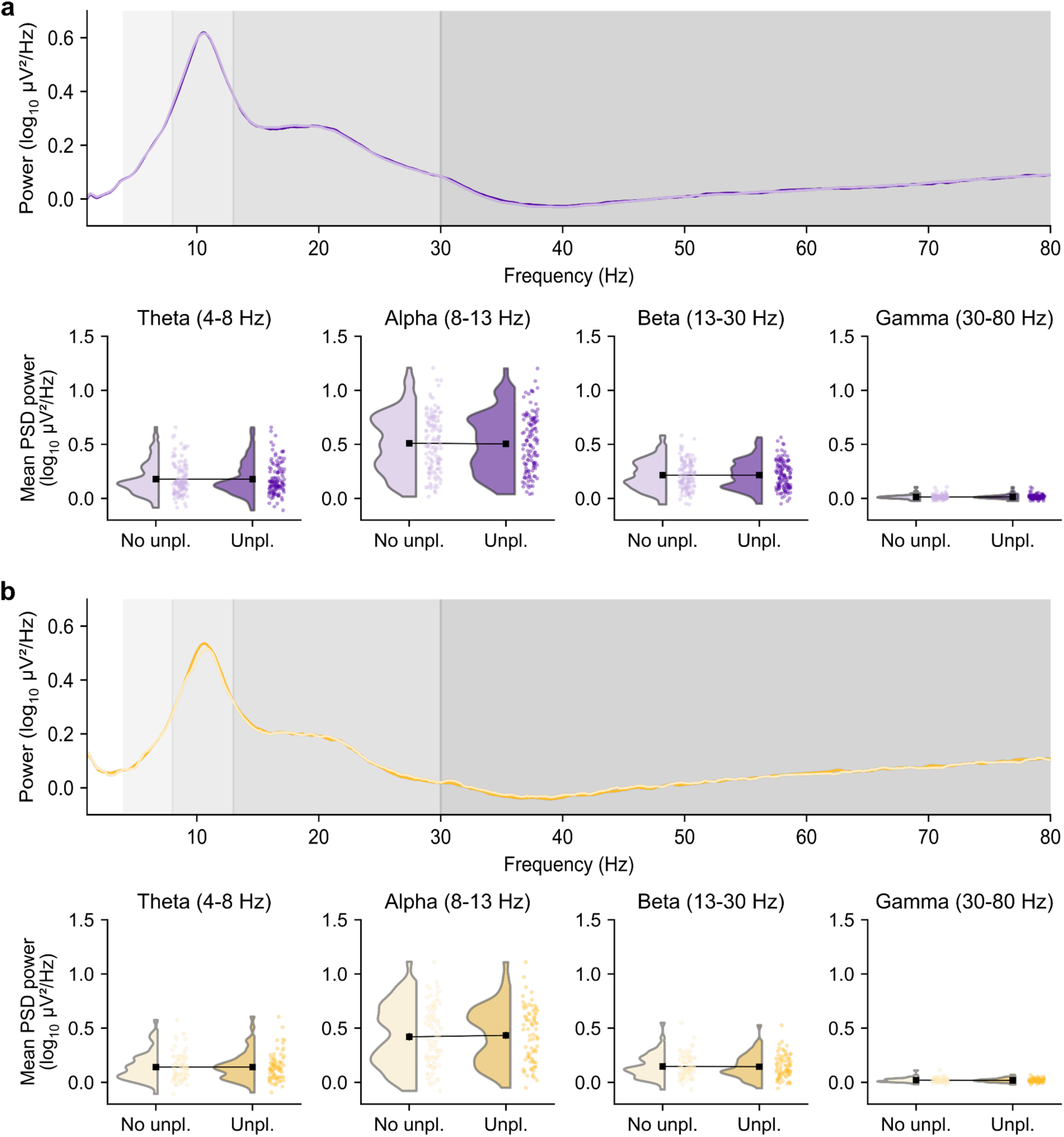
Global periodic PSD based on subjective ratings for aversive (a) auditory and (b) taste tasks. Lighter colors represent non-unpleasant epochs (VAS ≤ 99), while darker colors represent unpleasant epochs (VAS ≥ 100). Top panels display grand-average periodic power spectra across a frequency range of 4 to 80 Hz. Grey-shaded background regions indicate standard frequency bands: Theta (4–8 Hz), Alpha (8–13 Hz), Beta (13–30 Hz), and Gamma (30–80 Hz), respectively. Red horizontal bars show significant frequency clusters identified by the non-parametric cluster-based permutation tests (*p* < .05). The PSD between 58 to 62 Hz was interpolated to correct for the notch filter artifact. Bottom panels show the mean periodic PSD power by frequency band grouped by condition (no unpleasantness vs unpleasantness). Data are illustrated as raincloud plots, where the shaded violin plots represent the distribution of PSD power and individual dots represent mean PSD power for each participant. The black squares represent the group means. Unpl.: unpleasantness.

The spatial distribution of these modulations is visualized through topographical maps (Figures 7 and 8). For thermal pain stimulation (Figure 7a), the *t*-value map shows lower alpha and beta power during pain epochs compared to non-painful ones, particularly over the central and parietal regions, as well as higher gamma power over the frontal region. For chemical pain stimulation (Figure 7b), the difference maps revealed a pattern of increased power in the alpha band over centro-parietal regions during painful epochs. For both aversive tasks (Figure 8), overall *t*-value magnitudes were lower compared to pain stimulations. A mixed pattern of local increases and decreases was observed in all frequency bands for both tasks.

**Figure 7.**
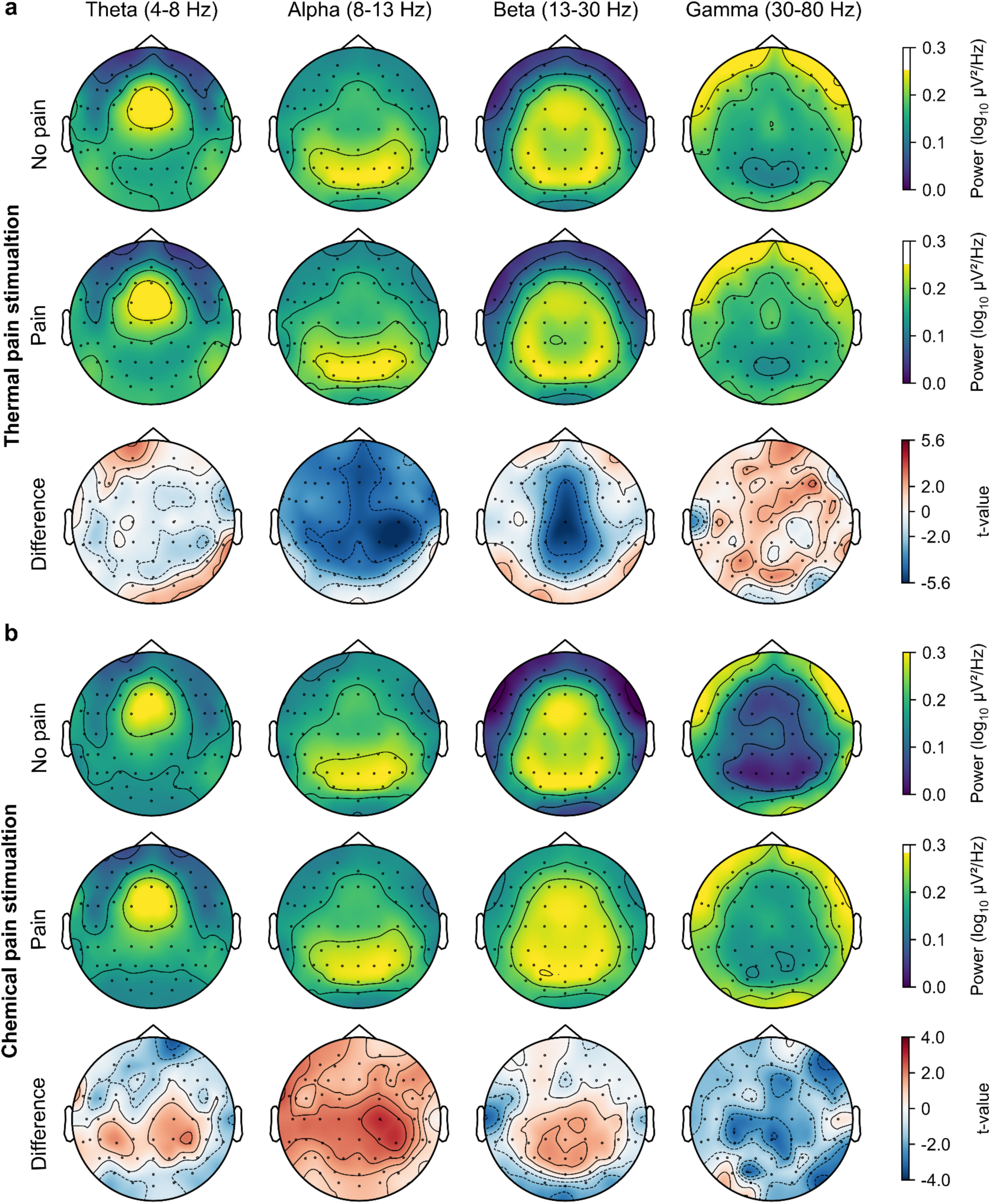
Global EEG topographical maps based on subjective ratings for (a) thermal pain and (b) chemical pain tasks. Maps are displayed for four frequency bands: theta, alpha, beta, and gamma. For each task, rows depict spatial distributions for no pain ratings (top), pain ratings (middle), and the resulting statistical *t*-value map of the difference (bottom). The critical *t*-value for statistical significance (*p* < .05) is represented on the color bars (i.e., ± 1.99).

**Figure 8.**
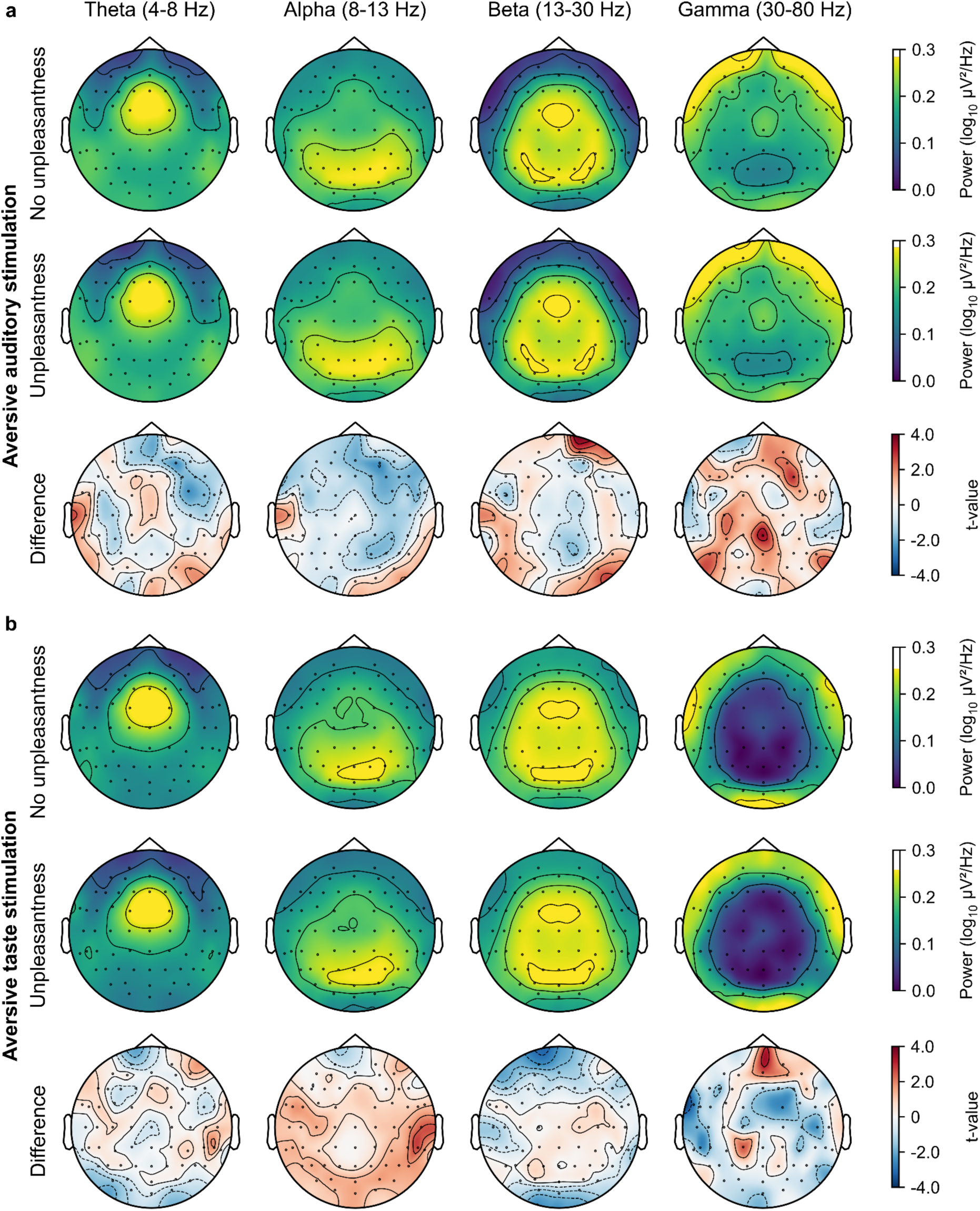
Global EEG topographical maps based on subjective ratings for aversive (a) auditory and (b) taste tasks. Maps are displayed for four frequency bands: theta, alpha, beta, and gamma. For each task, rows depict spatial distributions for no unpleasantness ratings (top), unpleasantness ratings (middle), and the resulting statistical *t*-value map of the difference (bottom). The critical *t*-value for statistical significance (*p* < .05) is represented on the color bars (i.e., ± 1.98 or 1.99).

**Figure 9.**
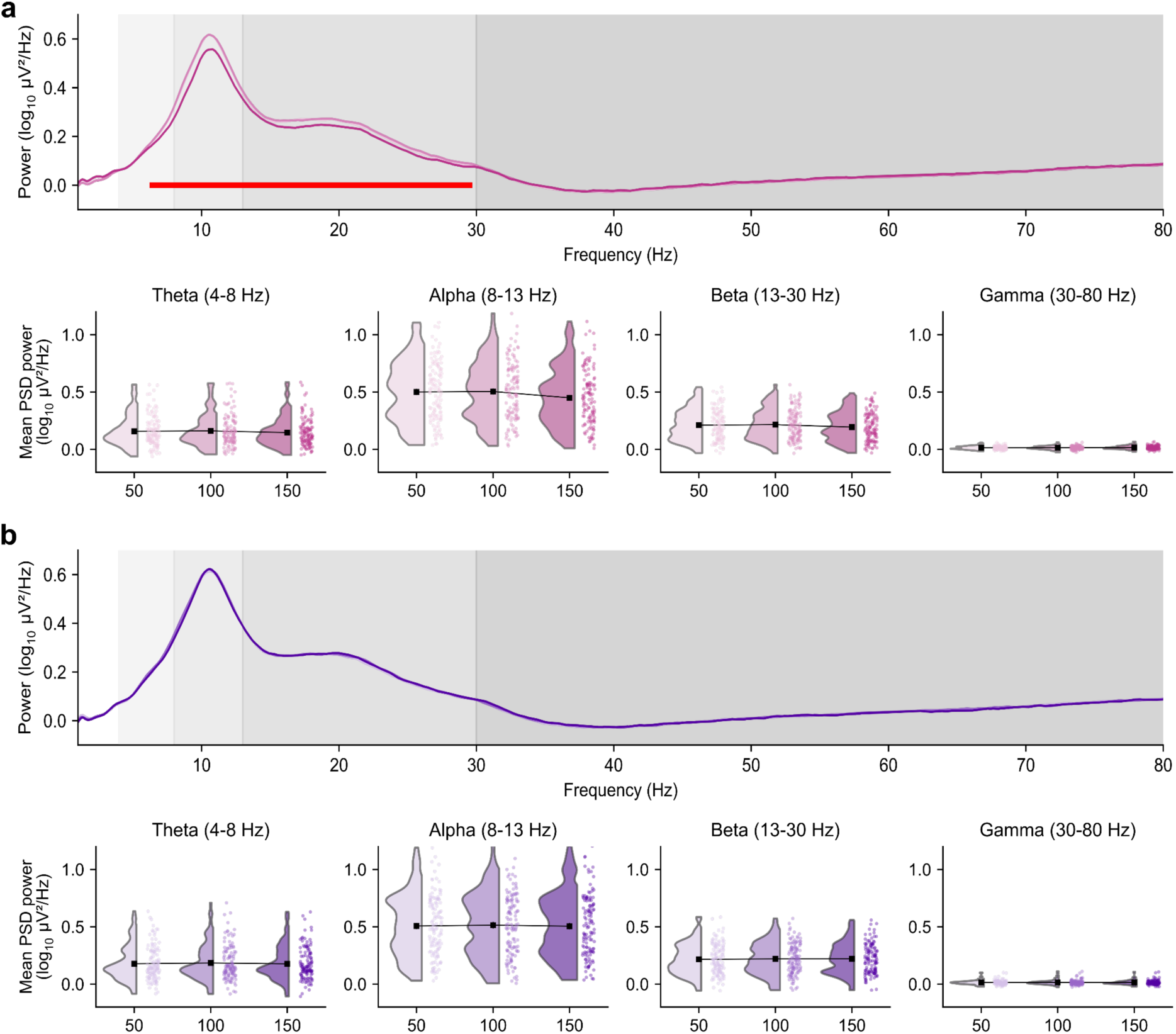
Global periodic PSD based on stimulus intensity for (a) thermal pain and (b) aversive auditory tasks. Top panels display grand-average periodic power spectra across a frequency range of 4 to 80 Hz. Grey-shaded background regions indicate standard frequency bands: Theta (4–8 Hz), Alpha (8–13 Hz), Beta (13–30 Hz), and Gamma (30–80 Hz), respectively. Red horizontal bars show significant frequency clusters identified by the non-parametric cluster-based permutation tests (*p* < .05). The PSD between 58 to 62 Hz was interpolated to correct for the notch filter artifact. Bottom panels show the mean periodic PSD power by frequency band grouped by target intensity (VAS-50, VAS-100, VAS-150). Data are illustrated as raincloud plots, where the shaded violin plots represent the distribution of PSD power and individual dots represent mean PSD power for each participant. The black squares represent the group means. 50: Target intensity VAS-50 (lightest color), 100: target intensity VAS-100 (medium color), target intensity VAS-150 (darkest color).

**Figure 10.**
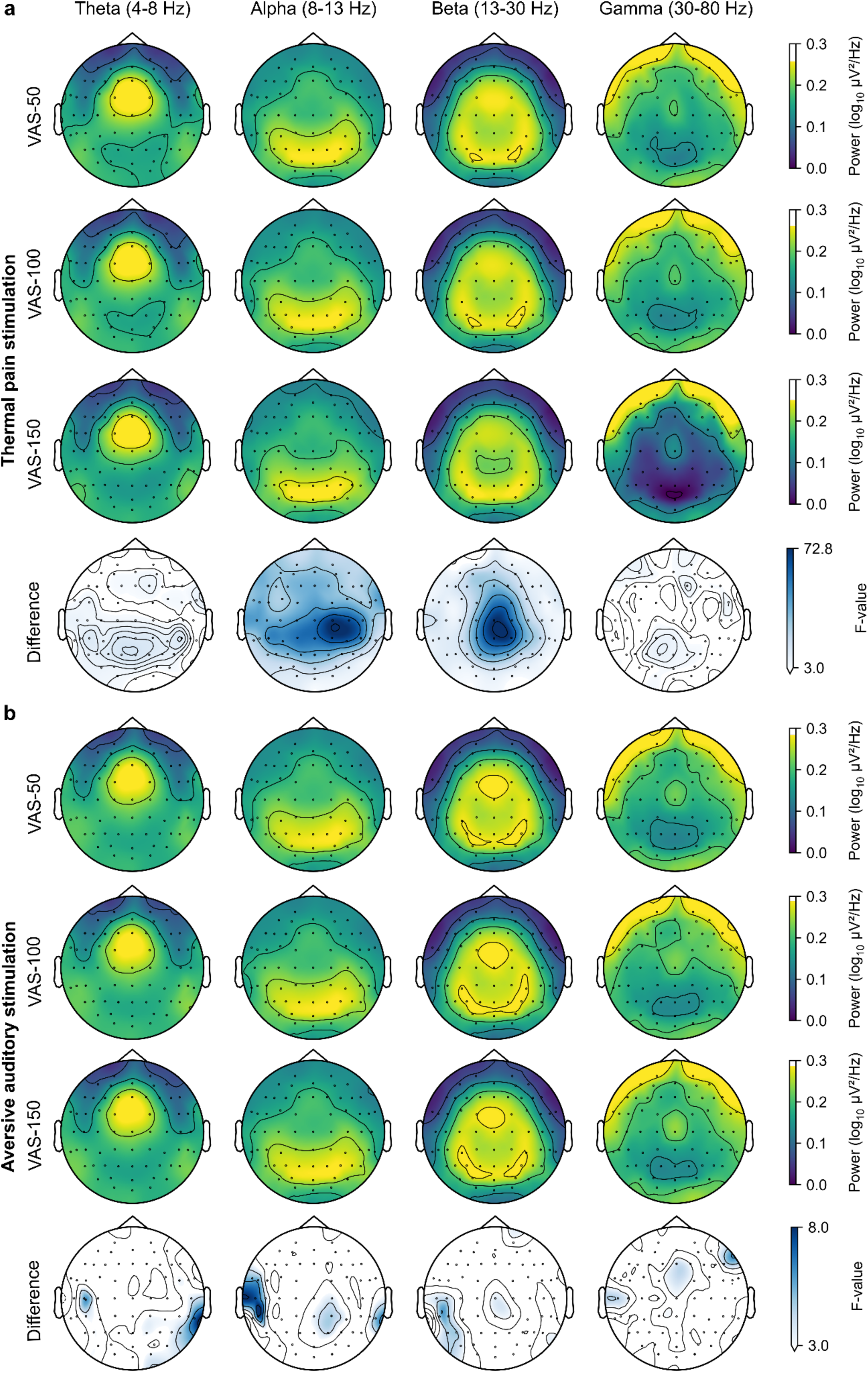
Global EEG topographical maps of stimulations intensity for (a) the thermal pain and (b) the aversive auditory tasks. Maps are displayed for four frequency bands: theta, alpha, beta, and gamma. For each task, rows depict spatial distributions for the three target intensities: VAS-50, VAS-100, and VAS-150, respectively. The fourth row shows the resulting *F*-value map of the difference between all three target intensities. The critical *F*-value for statistical significance (*p* < .05) is represented on the color bars (i.e., 3.02).

We first verified whether the EEG signal could differentiate between subjective states of “no pain/unpleasantness” and “pain/unpleasantness”. In the thermal pain task, non-parametric cluster-based permutation tests identified significantly lower power during painful rather than non-painful epochs in the theta, alpha and beta bands (indicated by red horizontal bars in Figure 5a). No significant frequency clusters were found in the chemical pain (Figure 5b), aversive auditory and taste tasks (Figure 6).

We then evaluated the sensitivity of the EEG data to variations in stimulus intensity for both the thermal pain and aversive auditory tasks (Figure 9 and 10). For the thermal pain target intensities, non-parametric cluster-based permutation tests identified significant frequency clusters in the theta, alpha, and beta bands, and none in the gamma band (see red horizontal bars in Figure 9a). The *F*-value map indicates a difference in alpha and beta power over the centroparietal regions (Figure 10a). For the aversive auditory target intensities, no significant frequency clusters were found (Figure 9b) and few significant differences in spatial distribution were observed (Figure 10b).

In summary, these topographical and spectral results indicate that our EEG data varies both according to the manipulation of stimulus intensity and the subjective ratings provided by participants. Notably, thermal pain showed a consistent pattern of modulations predominantly in the alpha and beta bands over central and parietal regions. Furthermore, the presence of modulations across all tasks supports the dataset’s validity for investigating neurophysiological changes associated with various painful and aversive modalities. While these initial findings likely capture confounding factors, such as motor activity related to continuous ratings, more sophisticated analytical approaches on this dataset will be able to disentangle these variables from specific pain-related activity.

### GSR and ECG responses

To validate the biological plausibility of the autonomic signals, we assessed the distribution and sensitivity of skin conductance and cardiac metrics across all experimental tasks. We also examined the sensitivity of these signals to subjective pain/unpleasantness perception, and to stimulus intensity for the aversive auditory and thermal pain tasks. GSR and ECG data based on subjective perception and on stimulus intensity are presented in Figure 11 and 12, respectively. Statistical results based on subjective perception are presented in Table 3, and those based on stimulus intensity in Table 4. For the auditory and thermal tasks, SCR magnitude reflected the first response exceeding the 0.01 μS threshold with its onset within a 5-8 s window within each intensity-specific epoch, and ΔHR was computed over a 20 s plateau from 4 to 24 s. Since intensity was not manipulated for the taste and chemical tasks, physiological metrics for the subjective analyses reflect only the first 20 s of stimulation.

**Figure 11.**
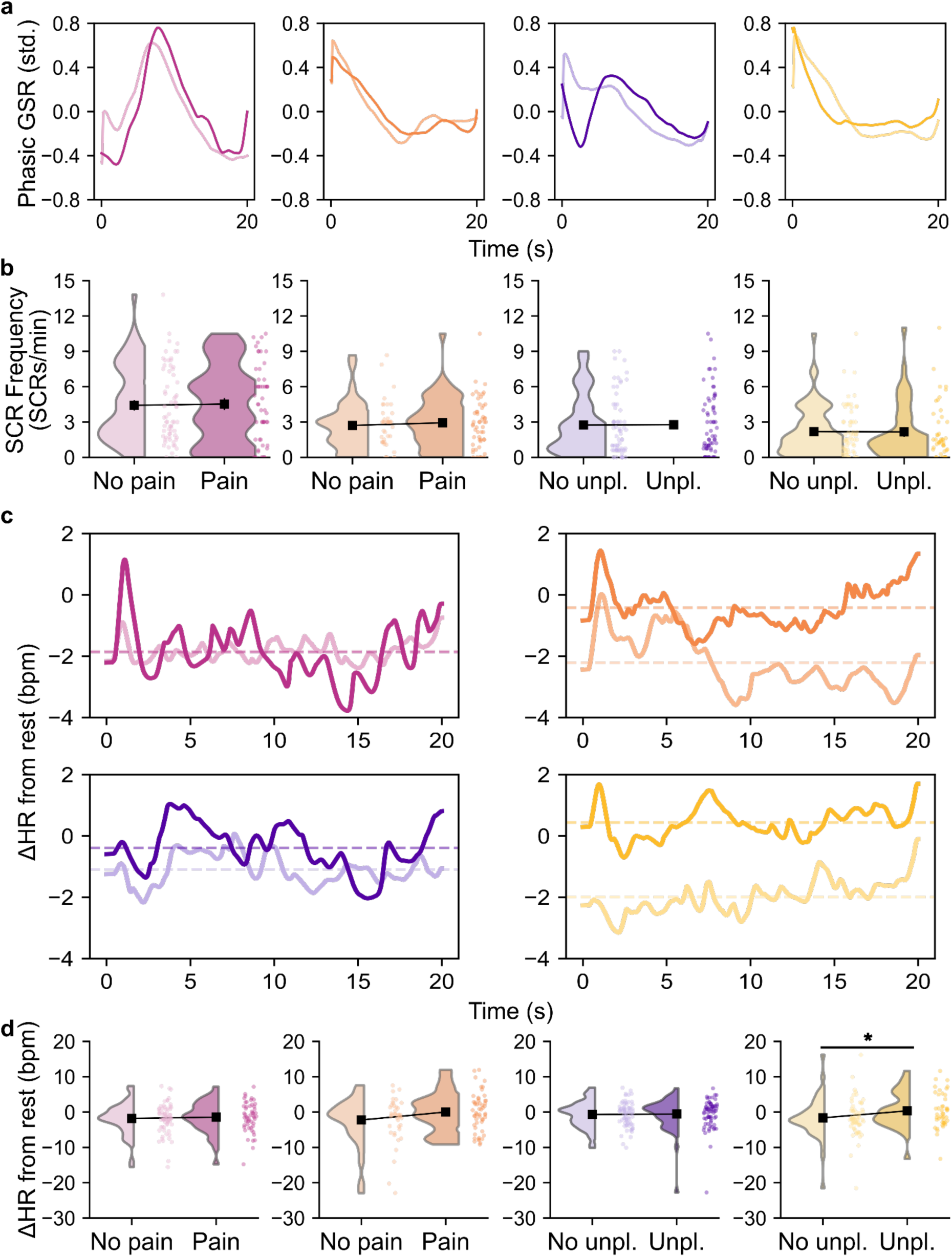
Physiological data based on subjective ratings. Each task is represented by a color: pink = thermal pain stimulation, orange = chemical pain stimulation, purple = aversive auditory stimulation, yellow = aversive taste stimulation. For all tasks, subjective ratings of pain or unpleasantness (Unpl.) correspond to the darkest shade, while ratings of no pain or no unpleasantness are shown with the lightest shade. Line plots display the grand average signal over time; in (c), dashed lines indicate the mean ΔHR value for each condition. Raincloud plots display the data distribution (half-violin), individual subjects means (points), and group mean (black square). Black horizontal lines with an asterisk (*) indicate statistically significant differences between conditions (*p* < .05). **(a) Standardized phasic GSR curve.** Grand average of standardized phasic GSR curves over 20 s epochs time-locked to stimulus onset (auditory and thermal), or over successive 20 s epochs spanning the recording excluding the first 10 s of stimulation (taste and chemical). **(b) SCR frequency.** Distribution of SCR frequency during stimulation (SCRs/min) per condition. **(c) ΔHR from rest curve.** Grand average of ΔHR from resting-state baseline curves over 20 s epochs beginning 4 s post-stimulus onset (auditory and thermal), or over the same successive 20 s epochs described in (a) (taste and chemical). **(d) ΔHR from rest.** Distribution of mean ΔHR from resting-state baseline (bpm) per condition. GSR: galvanic skin response; SCR: skin conductance response (frequency, SCRs/min); ΔHR: change in heart rate from resting-state baseline (bpm).

**Figure 12.**
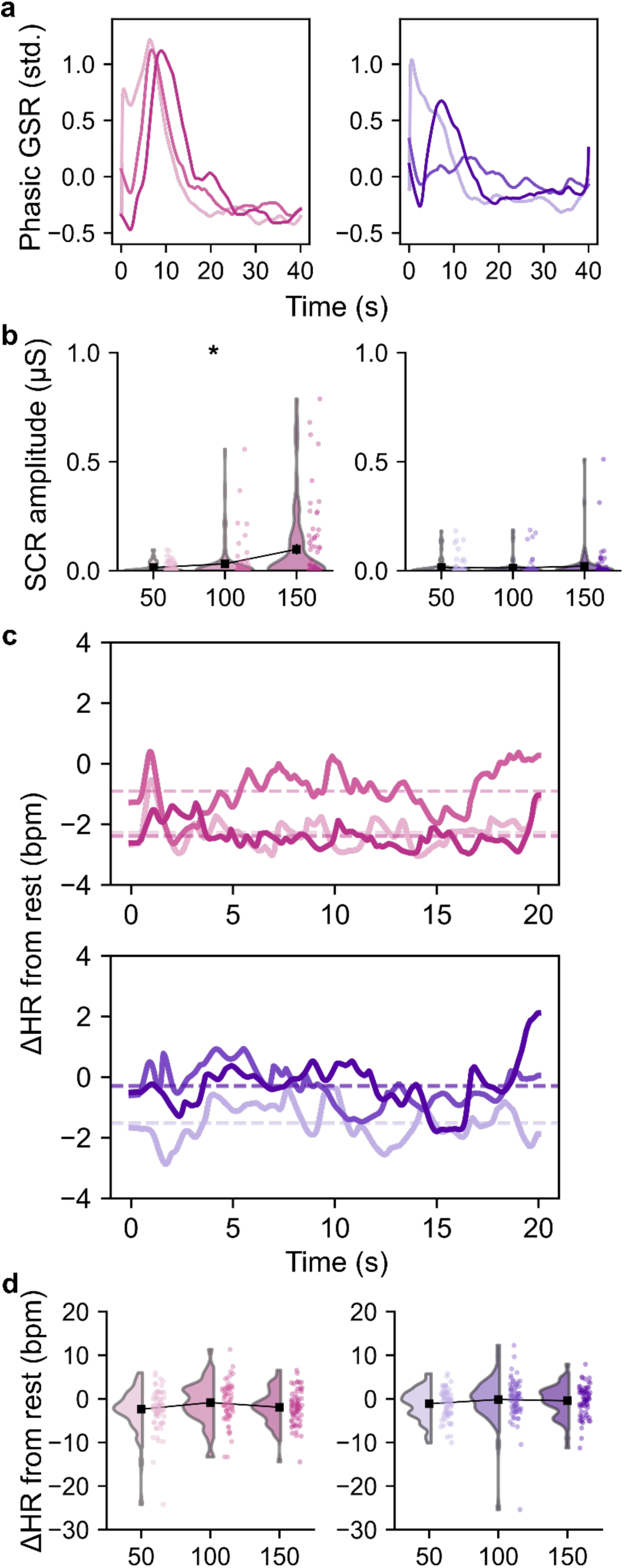
Physiological data based on stimulus intensity. Pink plots represent the thermal pain task, while purple plots represent the aversive auditory task. Target intensities are represented by the color shade: lightest = VAS-50 (50), intermediate = VAS-100 (100), darkest = VAS-150 (150). Line plots display data distribution over time, and dashed lines represent the mean value for each condition. Raincloud plots display the data distribution (half-violin), individual subjects means (points), and group mean (black square). An asterisk (*) represent the presence of a main effect of stimulus intensity based on LMM results (see Table 4). **(a) Standardized phasic GSR curve.** Grand average of standardized phasic GSR curves over 40 s epochs time-locked to intensity change onset. **(b) SCR amplitude.** Distribution of mean SCR amplitude per intensity condition. **(c) ΔHR from rest curve.** Grand average of ΔHR from resting-state baseline curves over 20 s epochs beginning 4 s post-stimulus onset. **(d) ΔHR from rest.** Distribution of mean ΔHR from resting-state baseline (bpm) per condition. GSR: galvanic skin response; SCR: skin conductance response; HR: heart rate.

**Table 3.** Statistical results of LMM for physiological metrics based on subjective experience across all experimental tasks. Fixed effects represent the contrast between painful/unpleasant (VAS ≥ 100) and non-painful/non-unpleasant (VAS ≤ 99) epochs. Beta (β) represents the fixed effect estimate in within-subject standardized units; SCR values were additionally log(1+x)-transformed prior to standardization. Bold p-values indicate significance after FDR correction (*p* < .05). Asterisks (*) denote models where the maximal structure was simplified to a random intercept only due to non-convergence of the model including both random intercept and slope. SCR: skin conductance response (frequency, SCRs/min); ΔHR: change in heart rate from resting-state baseline (bpm).

| Task | Metric | *n* | Beta ($\beta$) | *p* (FDR) | 95% CI |
| --- | --- | --- | --- | --- | --- |
| Thermal pain | SCR | 68 | 0.041 | .768 | [-0.10, 0.18] |
| | ΔHR* | 68 | 0.071 | .541 | [-0.07, 0.21] |
| Chemical pain | SCR | 67 | -0.094 | .541 | [-0.29, 0.10] |
| | ΔHR | 67 | -0.056 | .839 | [-0.38, 0.27] |
| Aversive auditory | SCR | 69 | -0.001 | .983 | [-0.12, 0.12] |
| | ΔHR* | 69 | 0.063 | .541 | [-0.06, 0.18] |
| Aversive taste | SCR | 69 | -0.079 | .541 | [-0.22, 0.06] |
| | ΔHR | 69 | 0.242 | **.034** | [0.08, 0.41] |

**Table 4.** Statistical results of LMM for physiological metrics based on stimulus intensity for the thermal pain and aversive auditory tasks. Fixed effect represents the linear trend across the three target intensities (VAS-50, VAS-100, and VAS-150), with intensity centered on VAS-100 and rescaled to unit steps of 50. Beta (β) represents the fixed effect estimate in standardized units per one-level increase in target intensity. SCR values were additionally log(1+x)-transformed prior to standardization. Bold *p*-values indicate significance after FDR correction (*p* < .05). Asterisks (*) denote models where the maximal structure was simplified to a random intercept only due to non-convergence of the model including both random intercept and slope. SCR: skin conductance response (magnitude of first detected response exceeding the 0.01 μS threshold with its onset within a 5-8 s window); ΔHR: change in heart rate from resting-state baseline.

| Task | Metric | *n* | Beta ($\beta$) | *p* (FDR) | 95% CI |
| --- | --- | --- | --- | --- | --- |
| Thermal pain | SCR* | 68 | 0.257 | **< .001** | [0.167, 0.347] |
|  | HR | 68 | 0.044 | .321 | [-0.029, 0.116] |
| Aversive auditory | SCR | 69 | 0.028 | .599 | [-0.075, 0.131] |
|  | HR* | 69 | 0.070 | .098 | [0.000, 0.138] |

Regarding electrodermal activity, we first confirmed that SCR frequency and SCR magnitude fell within expected physiological ranges^87^. Mean SCR frequency ranged from 2.2 to 4.5 responses/minutes across tasks, with non-response rates in 3 to 10% of epochs. After FDR correction, no significant effect of subjective experience was observed for SCR frequency in any task (Table 3). Mean SCR magnitude ranged from 0.013 to 0.098 μS across intensity conditions. However, the proportion of epochs in which no detectable SCR occurred was high in the objective intensity analyses, with 46% and 20% in the aversive auditory and thermal pain tasks, respectively. Since these analyses aimed to detect the first SCR and extract its magnitude, a recommended, smaller time window was used^47,69^. Individual variability in response latency or electrodermal lability^68,69^ is therefore more likely to result in a missed detection compared to the frequency-based measure. It is also possible that some intensity levels, as well as the decreasing intensity steps (see Figure 2), were insufficient to elicit a detectable SCR. After FDR correction, a significant positive linear trend was observed in the thermal pain task, indicating that higher target intensities were associated with larger SCR magnitude within participants (see Table 4). No significant intensity effect was found in the aversive auditory task.

Similarly, cardiac metrics were also biologically plausible, with mean absolute HR values ranging from 67.6 to 72.2 bpm across tasks. After FDR correction, a significant effect of subjective experience on ΔHR was observed in the aversive taste task (Table 3), where reporting unpleasantness was associated with higher ΔHR within participants. No significant effect of subjective experience on ΔHR was found in the aversive auditory or thermal pain tasks. Regarding stimulus intensity, a significant positive linear relation was observed in the aversive auditory task (Table 4), indicating that higher target intensities were associated with greater cardiac acceleration compared to rest. No significant intensity effect was found in the thermal pain task.

Taken together, these physiological patterns align with the current literature suggesting that GSR and HR are sensitive to heightened pain, arousal or negative emotions such as disgust^88,89^. The different patterns of sensitivity across metrics, where ΔHR responds more consistently to subjective experience and SCR amplitude reflects graded intensity changes in the thermal pain task, is consistent with prior findings that heart rate better predicts between-subject differences in pain ratings, whereas skin conductance is more sensitive to within-subject, intensity-related variation^19^. This is also compatible with the notion that electrodermal and cardiac responses capture partially dissociable aspects of autonomic arousal^89^. Furthermore, the absence of significant SCR frequency effects across all subjective analyses can be biologically plausible, as a frequency-based approach of non-specific phasic responses may be less sensitive to brief epoch-level fluctuations in perceived pain or unpleasantness than event-related amplitude measures^68^.

## Data Availability

The dataset is fully available on the following repository: https://openneuro.org/datasets/ds008349. All participants provided informed consent, explicitly agreeing to the public sharing of their data. Data sharing is conducted in accordance with ethical approval procedures, ensuring the protection of participants’ privacy.

## Code Availability

All latest versions of the code associated with this work have been made publicly accessible here: https://github.com/a-champagnee/eegmarkers_database.

## Author Contributions

A.C. conducted this research, analysed and interpreted the data, and wrote the manuscript. L.-E.B and A.C.-B. conducted this research. M.R. conceived this research. M.-P.C. acquired funding, conceived this research, provided support for data analysis and interpretation, and substantively revised this manuscript. All authors revised and reviewed the manuscript.

## Acknowledgments

A.C. and this work were supported by graduate scholarships from the Fonds de Recherche Québec - Nature et technologies (332150; https://doi.org/10.69777/332150) (345974; https://doi.org/10.69777/345974), the Natural Sciences and Engineering Research Council of Canada, and the Canadian Institutes of Health Research.

## Acknowledgements

M.-P.C. and this work were supported by a career grant and a start-up grant from the *Fonds de Recherche Québec -Santé* (309041), an operating grant from the *Fonds de Recherche Québec - Nature et technologies* (377430; https://doi.org/10.69777/377430) and the *Natural Sciences and Engineering Research Council of Canada* (ALLRP 608225 - 25), and a pilot project grant from the Quebec Bio-Imaging Network. A.C. and this work were supported by graduate scholarships from the *Fonds de Recherche Québec - Nature et technologies* (332150; https://doi.org/10.69777/332150) (345974; https://doi.org/10.69777/345974), the *Natural Sciences and Engineering Research Council of Canada*, the *Canadian Institutes of Health Research*, and a student grant from the *Canadian Pain Society*.

